# Somatic Mutations in Clonally Expanded T-lymphocytes in Patients with Chronic Graft-Versus-Host Disease

**DOI:** 10.1101/747196

**Authors:** Giljun Park, Daehong Kim, Jani Huuhtanen, Sofie Lundgren, Rajiv K. Khajuria, Ana M. Hurtado, Cecilia Muñoz-Calleja, Laura Cardeñoso, Valle Gómez-García de Soria, Tzu Hua Chen-Liang, Samuli Eldfors, Pekka Ellonen, Sari Hannula, Oscar Bruck, Anna Kreutzman, Urpu Salmenniemi, Tapio Lönnberg, Andres Jerez, Maija Itälä-Remes, Mikko A. I. Keränen, Satu Mustjoki

**Author notes:** **Corresponding author:** Prof. Satu Mustjoki, Hematology research unit Helsinki, University of Helsinki and Helsinki University Hospital Comprehensive Cancer Center, P.O. Box 700, Haartmaninkatu 8 FIN-00029 Helsinki, Finland.

## Abstract

Graft-versus-host-disease (GvHD) is the main complication of allogeneic hematopoietic stem cell transplantation. GvHD patients have aberrant T cell expansions, which are thought to drive pathological immune activation. Here we report mechanistic insights that somatic mutations may account for persistent clonal T cell expansions in chronic GvHD (cGvHD). In an index patient suffering from cGVHD, we discovered persisting somatic *MTOR*, *NFKB2*, and *TLR2* mutations in an expanded CD4+ T clone. In the screening cohort (n=135), the *MTOR P2229R* kinase domain mutation was detected in two additional cGvHD patients, but not in controls. Functional analysis of the discovered *MTOR* mutation indicated a gain-of-function alteration in translational regulation yielding in up-regulation of phosphorylated S6K1, S6, and AKT. Paired single-cell RNA and T cell receptor alpha and beta sequencing strongly supported cytotoxicity and abnormal proliferation of the clonally expanded CD4+ T cells. Real-time impedance measurements indicated increased cytotoxicity of mutated CD4 + T cells against the patient’s fibroblasts. High throughput drug-sensitivity testing suggested that mutations induce resistance to mTOR inhibitors but increase sensitivity for HSP90 inhibitors. Our findings suggest a novel explanation for the aberrant, persistent T cell activation in cGvHD, and pave the way for novel targeted therapies.

## INTRODUCTION

Graft-versus host disease (GvHD) is the main complication of allogeneic hematopoietic stem cell transplantation (allo-HSCT).(*1*) Chronic GvHD (cGvHD), that occurs more than 100 days after the transplantation, develops in 30-70 % of allo-HSCT recipients. Affected patients frequently need immunosuppressive treatment for years or even for a lifetime, and in many patients the condition is fatal.(*2*) The genesis of cGvHD is multifactorial, but donor alloreactive lymphocytes are believed to be the key pathogenetic drivers that target host tissues such as skin, soft tissues, oral mucosa, and eyes. In particular, CD4+ T cells contribute to the early inflammation and tissue injury, to subsequent chronic inflammation, and late aberrant tissue repair and fibrosis.(*3*)

During normal immune response, naïve T cells encounter their cognate antigen, get activated and undergo a rapid clonal expansion.(*4*) The activation and proliferation of T cells are usually tightly regulated processes, in which effector cells undergo apoptosis upon proper immune response. In many immune-system-mediated disorders, such as cGvHD, the immune homeostasis is disturbed, and the enormous variability of different T cell clones is diminished. In some patients, the T cell receptor (TCR) repertoire is heavily skewed, and clones comprising up to 20-40% of all T cells can exist.(*5*) The underlying mechanisms for this phenomenon remain unknown.

Here, we hypothesized that in cGvHD antigen-encountered T cells may acquire somatic mutations due to constant immune-system activation and proliferation. Such mutations might lead to functional and survival advantages of T cells that result in clonal expansion and aberrant immune responses. To explore this theory, we sequenced purified CD4+ and CD8+ lymphocytes from an index patient suffering from cGVHD with a custom deep-sequencing panel consisting of immunity and inflammation-related genes. Mutation findings were confirmed in a validation cohort of 135 GVHD patients. Subsequently, the functional consequences of the discovered mutations including their role in conferring resistance to immunosuppressive therapy were evaluated *in vitro*, and finally verified with patient cells using drug sensitivity screening and unbiased transcriptome-wide single-cell RNA-sequencing (scRNA-seq) paired with T cell receptor alpha and beta sequencing (TCRab-seq) analysis and functional cytotoxicity assays.

## RESULTS

### Clinical Characteristics of the Index cGvHD Patient

The index patient was a 56-year-old male, who was diagnosed with chronic phase chronic myeloid leukemia in 1999. The clinical status and treatment history are described in detail in Supplemental Figure S1 and Supplemental results. Since the beginning of 2001, the patient suffered from cGvHD affecting his liver, eyes, nails, and skin. The immunosuppression was continuously adjusted according to the clinical presentation of the cGvHD.

### Immunophenotype and clonal expansions of CD4+ and CD8+ T cells

During the first sampling in 2013, the patient was on mycophenolate mofetil therapy. T cell clonality was initially analyzed with a flow cytometry-based assay using a panel of TCR Vβ-specific antibodies. A large clonal Vβ20+ expansion was noted among CD4+ T cells. The Vβ20+ clone constituted approximately 50% of pure CD4+T cells and 60% CD4+CD8+ T cells (Figure 1A and 1B). Additionally, smaller clonal expansions (Vβ5.1 11.8%, Vβ7.1 21.0%, Vβ17 12.1%, and Vβ23 12.2%) were detected among CD8+ T cells (Figure 1B). To assess the clonality in more detail, FACS-sorted CD4+ Vβ20+, CD4+ Vβ20- and CD8+ T cells, obtained in 2015, were further analyzed by a TCRβ deep-sequencing assay(*6*), which confirmed the TCRBV30-01 clone expansion (corresponding to the Vβ20+ expansion observed by flow cytometry) in the CD4+Vβ20+ fraction (66.2% of sorted cells) (Figure 1C). TCRβ sequencing of CD8+ T cells revealed two relatively large clones, TCRBV07-09 (16.1%) and TCRBV28-01 (17.9%).

**Figure 1.**
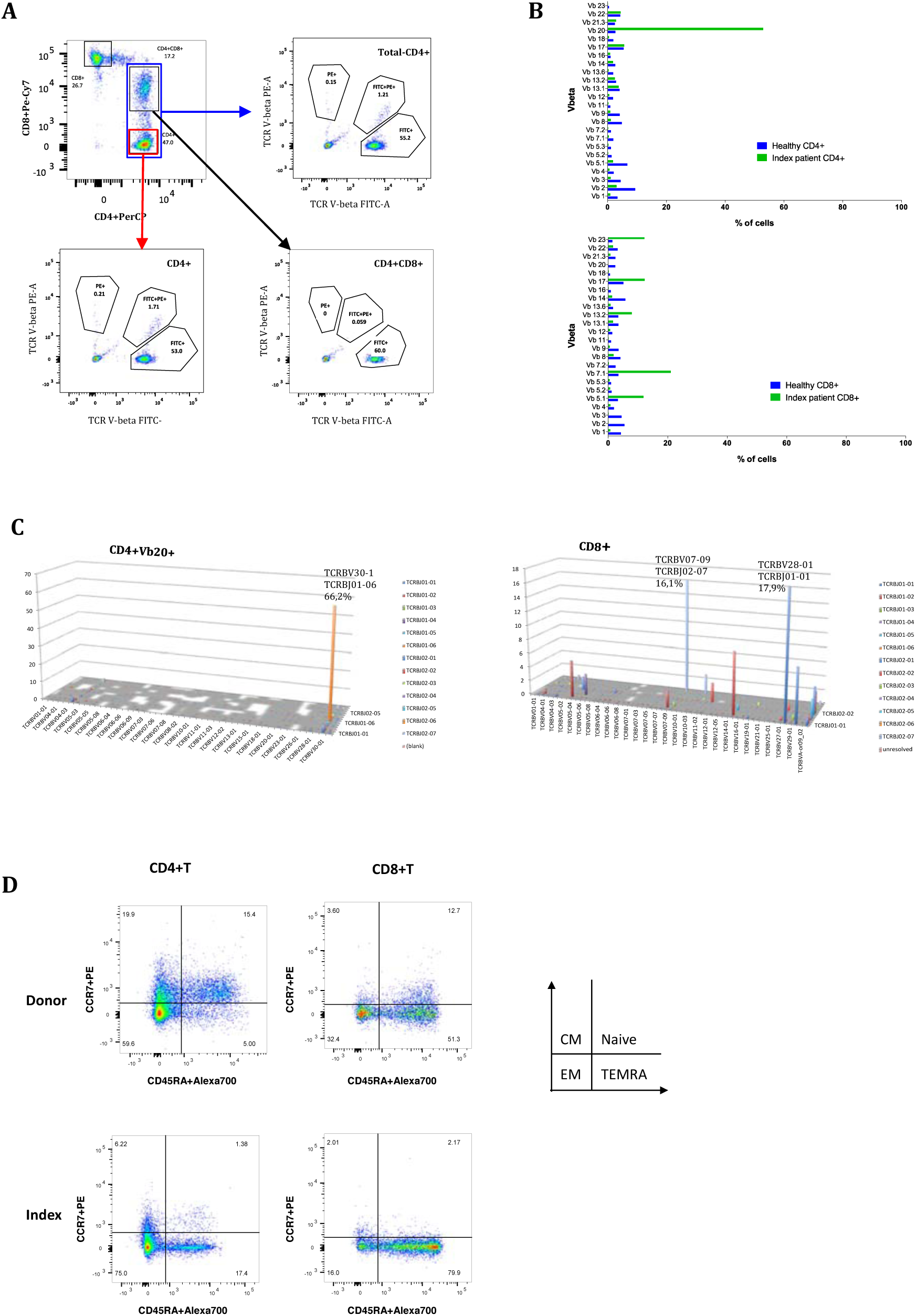
Flow cytometry and TCRB deep sequencing results from the index patient. (A) and (B). TCR Vβ repertoire of CD4+ and CD8+ T cells was analyzed in peripheral blood from the Index patient with the IO Test Beta Mark TCR beta Repertoire Kit (Beckman-Coulter Immunotech, USA). 53% of CD4+T cells and 60% of CD4+CD8+T cells consisted of a single Vβ20 clone. (C) T cell repertoire of FACS-sorted CD4+Vβ20+ and CD8+ T cells analysed with TCRβ deep sequencing (Adaptive Biotech., USA). The TCRBV30-01 clone was detected in the CD4+Vβ20+ fraction, but not in the CD8+ fraction. (D) Multicolor flow cytometry was applied to identify the immune phenotype of donor and index patient’s memory T cell subtypes. Central memory (CM), naïve, effector memory (EM) and terminal effector memory (TEMRA) cells.

During an exacerbation of sclerodermatous skin lesions in 2015, 59% of peripheral blood leukocytes were T cells, 5% B cells, and 35% NK cells (Figure S2A in the Supplementary Appendix). CD3+ T cells were composed of CD4+ (47%), CD4+CD8+ (17%), and CD8+ T cells (27%) (Figure S2 in the Supplementary Appendix). An increased number of CD4+ effector memory (EM, 75.0%) and terminally differentiated effector memory (TEMRA) cells (17.4%) was found together with a decreased number of CD4+ central memory (CM) cells (6.2%) when compared with the sibling donor’s CD4+ T cell pool (59.6% EM, 5.0% TEMRA, and 19.9% CM cells)(Figure 1D). In the CD8+ T cell pool, increased amount of TEMRA cells was noted (79.9% of CD8+ T cells).

### Somatic Mutations in the Expanded CD4+ T Cell Population of the Index Patient

To screen for somatic mutations, a customized immunity and inflammation-related gene sequencing panel (immunogene panel)(*6*) was applied to immunomagnetic bead-separated blood CD4+ and CD8+ T cells, that were obtained from the index patient in 2013. The median target gene coverage for the panel was 152 in CD4+ and 160 for CD8+ T cells. In total, 14 candidate putative somatic mutations were discovered within the CD4+ T cells (Table 1), and one in CD8+ cells (Table S7A in the Supplementary Appendix). Based on the known biological significance, three of the mutations (*MTOR*, *NFkB2*, and *TLR2)* were considered as putative driver mutations and potentially important for disease pathogenesis and were studied further.

**Table 1.**
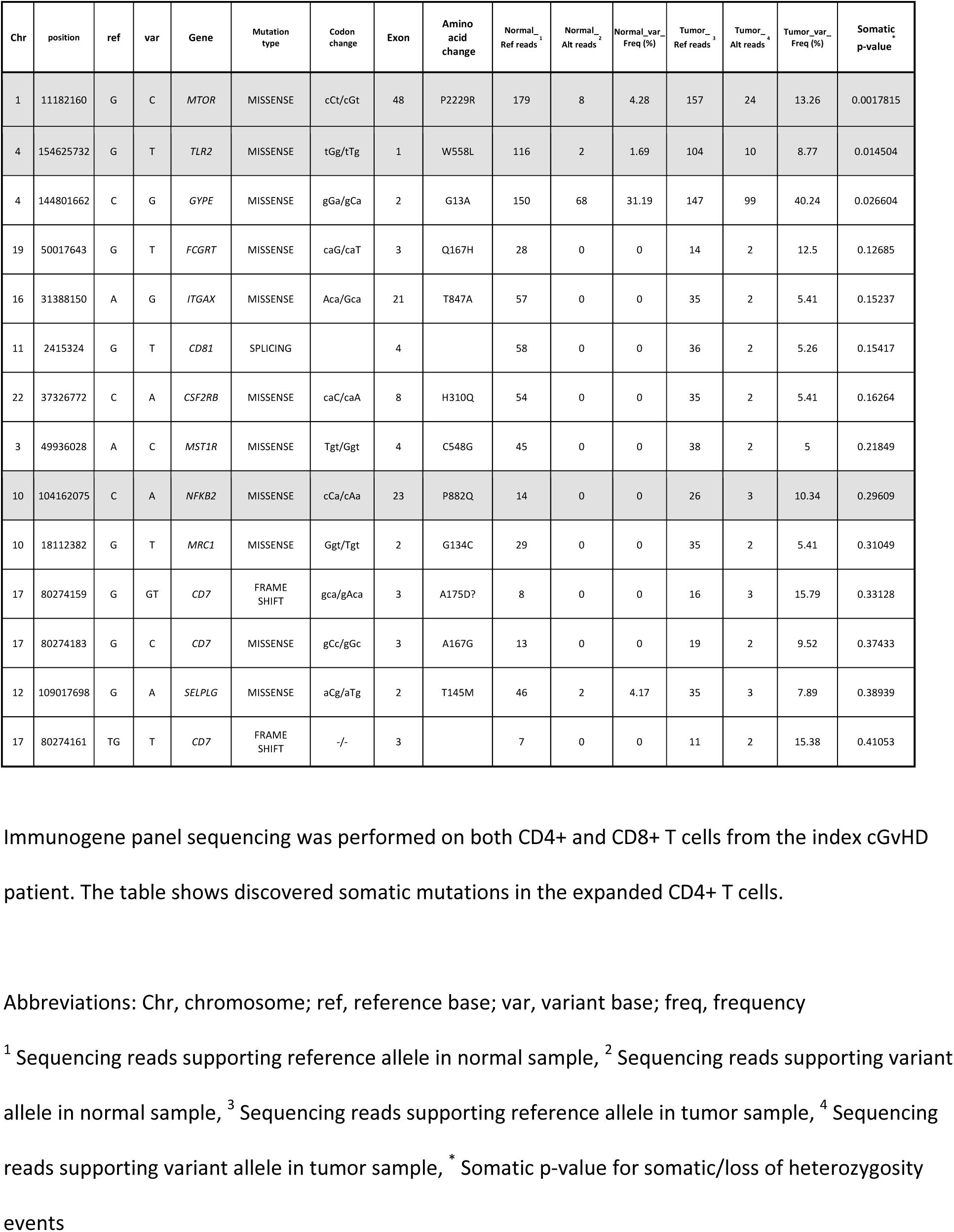
Somatic mutations discovered in CD4+ T cells in the index patient, detected from 2013 sample

The previously undescribed somatic missense mutation in *MTOR* (position 11182160, G to C) changes the amino acid proline 2229 to arginine (Figure 2A). The variant allele frequency (VAF) was 13.3% among CD4+T cells (Table 1). This mutation in exon 48 is located in the kinase domain which has been suggested to be important for signal transduction.(*7*) In addition to *MTOR*, two other interesting mutations were identified in the *NFkB2* and *TLR2* genes, although these were statistically not significant (p>0.01) due to the low coverage in these locations (Table 1). The *NFkB2* missense mutation (position 104162075, C to A) leads to a change of the amino acid proline 882 to glutamine (Figure 2A). *TLR2* missense mutation (position 154625732, G to T) results in a change of the amino acid tryptophan 558 to leucine (Figure 2A). The transcription factor NFkB2 is a critical regulator of inflammation and immune function.(*8*) Toll-like-receptor 2 (TLR2) is one of the pattern recognition receptors and has been shown to be a crucial player for the pathogenesis of autoimmune diseases. Notably, TLR2 protein has been shown to be highly expressed in GvHD patients.(*9*)

**Figure 2.**
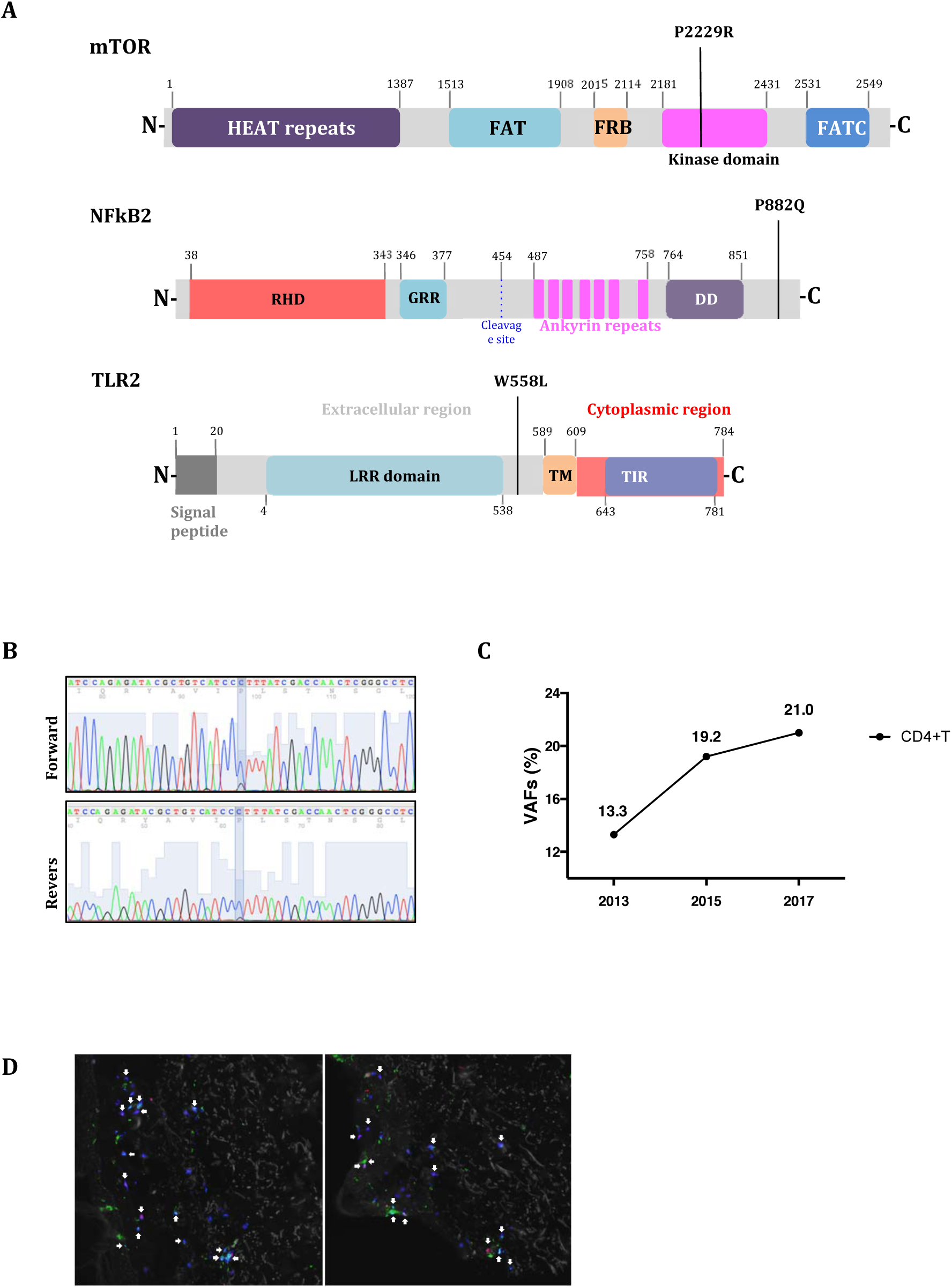
*MTOR, NFkB2*, and *TLR2* mutations in index patient. (A) Locations of *MTOR, TLR2*, and *NFkB2* somatic mutations. Linearized structure of MTOR, NFkB2, and TLR2 presenting the location of somatic mutations. *MTOR P2229R* mutation is located in the kinase domain, *NFkB2 P882Q* in the C-terminus, and *TLR2 W558L* between LRR (Leucine-rich repeats) domain and transmembrane (TM) domain. (B) A heterozygous *MTOR* mutation (G to C, P2229R) was detected in CD4+ T cells by Sanger sequencing. (C) Variant allele frequencies (VAFs) of *MTOR* mutation in the index patient’s CD4+ T cells over time as measured with amplicon sequencing. (D) Immunofluorescence staining indicated CD3+CD4+ and CD3+CD8+T cell infiltration in the skin. Paraffin embedded skin biopsy from index patient was sectioned and stained with antibody specific human CD3 (cyan), CD4 (green), and CD8 (red). White arrows indicate infiltrated CD3+CD4+ or CD3+CD8+T cells.

As an additional confirmation of these findings, we performed exome sequencing of CD4+, CD8+ T cells, and NK-cells obtained from the index patient in 2015 (Tables S7B, C, and D in the Supplementary Appendix). Altogether, 15 candidate putative somatic mutations were discovered within the CD4+ T cell population, including those in the *MTOR*, *TLR*2, and *NFkB2* genes (Table S7 in the Supplementary Appendix).

### Validation of the MTOR, TLR2, and NFkB2 Mutations in the Index Patient

To further validate the *MTOR P2229R*, *TLR2 W558L*, and *NFkB2 P882Q* mutations, CD4+ cells obtained in 2015 underwent standard capillary sequencing. Only the *MTOR* mutation was confirmed due to the low sensitivity of the assay (Figure 2B). Therefore, targeted amplicon sequencing with a coverage up to 100.000X and a sensitivity of 0.5% VAF(*10*) was applied to all available samples from different time points to establish the dynamics and the lineage specificity of the discovered mutations. The *MTOR* mutation was detected in CD4+ T cells that were obtained in 2015 and 2017 with VAFs 19.2 % and 21.0 %, respectively (Table 2 and Table S7 in the Supplementary Appendix). The VAF of the *MTOR* mutation appeared to increase from 2013 to 2017 regardless of the continuous immunosuppressive therapy (Figure 2C). No mutations or very low VAFs were detected within flow-sorted CD3-, CD8+/CD4-, CD4+Vβ20-, CD4+CD8+ Vβ20-, and monocyte samples (Figure S2C and Table S8 in the Supplementary Appendix). Thus, the mutation was confined to the CD4+ Vβ20+ cell fraction (VAF 44.7% in flow-sorted cells).

**Table 2.** Somatic *MTOR* and *NFkB2* mutations validated by amplicon sequencing

| Sample year | Patient | DNA | Gene | Chr | position | ref | var | Amino acid change | Call_Depth | Ref_Calls | Var_Calls | VAF (%) | Freq_Ratio |
| --- | --- | --- | --- | --- | --- | --- | --- | --- | --- | --- | --- | --- | --- |
| 2015 | 1-1 | CD4+ T | *MTOR* | 1 | 112182160 | G | C | P2229R | 1000004 | 808220 | 191784 | **19.2** | 1.00047 |
| 2015 | 1-2 | CD8+T | *MTOR* | 1 | 112182160 | G | C | P2229R | 1000003 | 942683 | 57320 | **5.7** | 0.9993 |
| 2017 | 1-3 | CD4+ Vb.20+ | *MTOR* | 1 | 112182160 | G | C | P2229R | 168592 | 93183 | 75409 | **44.7** | 0.99928 |
| 2017 | 1-4 | CD4+CD8+ Vb.20+ | *MTOR* | 1 | 112182160 | G | C | P2229R | 171728 | 110670 | 61058 | **35.5** | 0.99910 |
| 2015 | 1-5 | Skin-biopsy | *MTOR* | 1 | 112182160 | G | C | P2229R | 8085 | 7986 | 99 | **0.8** | 0.96305 |
| 2016 | 2 | Whole blood | *MTOR* | 1 | 112182160 | G | C | P2229R | 2743 | 2715 | 28 | **1.0** | 1.01793 |
| 2015 | 3 | Whole blood | *MTOR* | 1 | 112182160 | G | C | P2229R | 719 | 703 | 16 | **2.0** | 1.02671 |
| 2015 | 1-1 | CD4+T | *NFkB2* | 10 | 104162075 | C | A | P882Q | 471375 | 414821 | 56554 | **12.0** | 1.0061 |
| 2015 | 1-2 | CD8+T | *NFkB2* | 10 | 104162075 | C | A | P882Q | 586903 | 577283 | 9620 | **1.6** | 1.00124 |
| 2017 | 1-3 | CD4+ Vb.20+ | *NFkB2* | 10 | 104162075 | C | A | P882Q | 26262 | 20668 | 5594 | **21.3** | 1.00395 |
| 2017 | 1-4 | CD4+CD8+ Vb.20+ | *NFkB2* | 10 | 104162075 | C | A | P882Q | 48654 | 43822 | 4832 | **9.9** | 1.00282 |

CD4+ and CD8+ T cells were sorted either with the magnetic beads (2015 sample) or flow based sorting
(2017 sample). In addition, CD4+Vb20+ and CD4+CD8+Vb20+ fractions were sorted with flow cytometry
(2017 sample). *MTOR* and *NFkB2* mutations were analysed from sorted fractions with deep amplicon
sequencing. Mutations were confined to CD4+ fractions. The low mutation VAFs in CD8+ fraction are due
to small CD4+ T cell contamination (CD4+CD8+ double positive cells) in the bead sorted fraction.
Abbreviations: Chr, chromosome; ref, reference base; var, variant base; Call\_depth, total number of called
reads; Ref\_calls, sequencing reads supporting reference allele; Var\_calls, sequencing reads supporting
variant allele; VAF, variant allele frequency; Freq, frequency; Freq\_Ratio, base quality frequency ratio.

Similarly, the *NFkB2 P882Q* mutation was confirmed to be limited to CD4+Vβ20+ T cells (VAFs: 12.0% in CD4+ T cells and 21.3% in CD4+Vβ20+ cells) (Table 2, S7 and S8 in the Supplementary Appendix). In contrast, *TLR2 W558L* mutation was discovered in both CD4+ (VAFs: 16.5% in CD4+ cells, 35.5% in CD4+Vβ20+ cells) and CD8+ T cells (VAF 4.7 %) (Table S9 and S10 in Supplementary Appendix).

In the course of the disease, the cGvHD affected different organs of the index patient, but particularly the skin. To explore whether lymphocytes harboring the detected somatic mutations can be found in target organs, we screened paraffin-embedded biopsy samples by amplicon sequencing. The *MTOR* mutation was identified in a sclerodermatous lesion that was biopsied in 2015 (VAF 0.8 %), but not in eye or liver biopsies (Table 2). Immunofluorescence staining demonstrated CD4+ and CD8+ T cell infiltration in the same sclerodermatous lesion (Figure 2D).

To examine whether the mutations were already present in the donor, both CD4+ and CD8+ T cells from the donor were sequenced by amplicon sequencing, but no mutations were detected.

### Screening for the Identified *MTOR*, *TLR2*, and *NFkB2* Mutations in cGvHD patient Cohort

To explore whether the found mutations are recurrent, blood samples from 135 cGvHD patients, 38 allo-HSCT patients without cGvHD, and 54 healthy controls were screened by the amplicon sequencing. Two additional cGvHD patients carried the same *MTOR* missense mutation yielding in a *MTOR P2229R* mutation frequency of 2.2 % in all cGvHD patients (3 out of 135). In healthy controls or allo-HSCT patients without cGvHD, no *MTOR* mutations were detected. Furthermore, *NFkB2* mutations were neither detected in additional cGvHD patients nor in controls. The *TLR2 W558L* mutation was found in both cGvHD patients and healthy controls, but the VAF indicated a 10- and 4-fold higher mutation frequency in cGvHD patients’ CD4+ and CD8+ T cells compared to healthy controls (Figure S3, Tables S9 and S10 in the Supplementary Appendix).

### Identified Somatic Mutations Result in a Gain-of-function Alteration

MTOR consists of two functionally distinct multi-protein complexes, mTORC1 and mTORC2. Eukaryotic translation initiation factor 4E (eIF4E)-binding protein 1 (4E-BP1) and ribosomal S6 kinase (S6K1) are among the key substrates of mTORC1, and as such critical regulators of cap-dependent translation.(*11*) mTORC2 directly phosphorylates AKT, thereby promoting cell survival.(*12*) Various genomic alterations have been shown to aberrantly activate the mTOR pathways(*13*), which is marked by an increased phosphorylation of downstream factors, such as S6K1, S6, and AKT. To examine the functional consequences of the *MTOR P2229R* and other mutations, we transduced HEK293 human embryonic kidney cells with mutant constructs. Both the *MTOR P2229R* single mutant and triple mutant resulted in a substantially enhanced phosphorylation of S6K1, S6, and AKT compared to wild-type (WT) MTOR or triple WT (Figures 3A and B) suggesting an activation of both mTORC1 and mTORC2 pathways.

**Figure 3.**
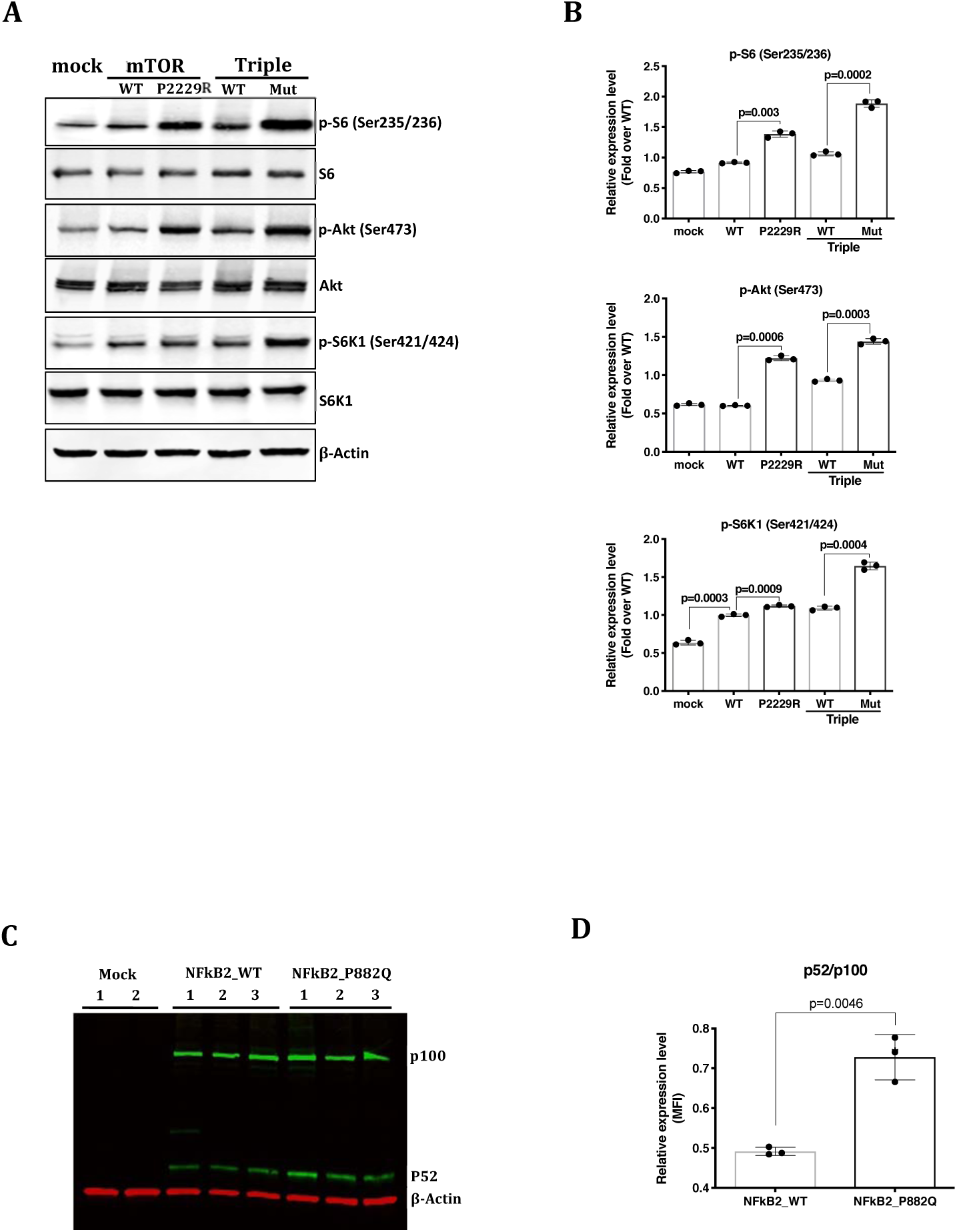
Functional analysis for wild type, *MTOR*, and *NFkB2* mutants in HEK293 cells. (A) Stably expressed *MTOR* wildtype (WT), mutant (P2229R), Triple (*MTOR, TLR2*, and *NFKB2*) wildtype (WT) and Triple mutant (*MTOR P2229R*, *TLR2 W559L*, and *NFKB2 P882Q*) (Mut) in HEK293 cells were serum starved for 12 hours. Western blot analysis was performed with the use of anti-pS6, anti-S6, anti-pAkt, anti-Akt, anti-pS6K1, anti-S6K1, and anti-β-actin. Data is representative of three independent experiments. (B) Relative expression level was estimated by measuring each band intensity of three independent experiments using ImageJ software (Rasband, W.S., ImageJ, U. S. National Institutes of Health, Bethesda, Maryland, USA, https://imagej.nih.gov/ij/, 1997-2016). Error bar present Mean ± SEM (n=3 per group). P values are derived from unpaired t-test with Welch’s correction (WT vs P2229R, WT vs Triple Mut, and mock vs WT in p-S6K1). (C) Immonoblot assay was performed to verify an alteration of P52 and P100 expression with *NFkB2* WT and *P882Q* mutant. Data is representative of three independent experiments. (D) Stably expressing *NFkB2 P882Q* increased expression of P52. Mean fluorescence intensity was measured by ImageJ software (NFkB2_WT: n =3, NFkB2_P882Q: n=3). Error bar present Mean ± SEM. P values are derived from unpaired t-test with Welch’s correction (NFkB2_WT vs NFkB2_P882Q).

The *NFkB2* P882Q mutation is located in the c-terminal domain (Figure 2A), which is known to play an important role in the ubiquitination and partial proteolysis from NFkB2 (p100) to NFkB2 (p52).(*14*) In order to determine the molecular balance between these two states, we performed immunoblotting which revealed an increased expression of p52 (Figures 3C and D) indicating a gain-of-function alteration.

Similarly, to study the functional consequences of the *TLR2 W558L* mutation, we evaluated alterations in transcriptional regulation by analyzing mRNA expression levels of a subset of *TLR2* downstream targets by qRT-PCR. This demonstrated significantly increased expression levels of *ELK1* and *FOS1* in the *TLR2 W558L* mutant expressing cell line as compared to the WT control (Figure S4B in the Supplementary Appendix).

### Paired scRNA- and TCRab sequencing of the CD4+ T cells from the index patient from two time points

Recently, entire transcriptome at the single-cell level has been explosively studied for revealing a differential gene expression profiles between individual cells, which cannot be identified from analysis of mixed cells.(*15*)

In order to better understand the heterogeneity of the clonotype and the underlying CD4+ T cell compartment in an unbiased manner, we performed simultaneous single-cell RNA and paired TCRab sequencing on two time points in 2015 and 2017 for the index patient’s CD4+ T lymphocytes from peripheral blood. From the paired sequencing we received 15,847 CD4+ T lymphocytes passing the quality control, and they could be divided into nine distinct phenotypes with graph-based clustering. Interestingly, most of the cells (72.0%) were characterized by cytotoxicity (clusters 0, 1, 2 and 3), and lower frequency of naïve cells (20.8%, clusters 4 and 5) and regulatory T cells (3.2%, cluster 8) were identified (Figure 4 A-C). The frequency of cells in clusters were stable between the time points indicating resistance to the ongoing immunosuppressive treatment, and only the two smallest populations showed over two-fold-change between the two timepoints (clusters 3 and 7, Figure S5 in the Supplementary Appendix).

**Figure 4.**
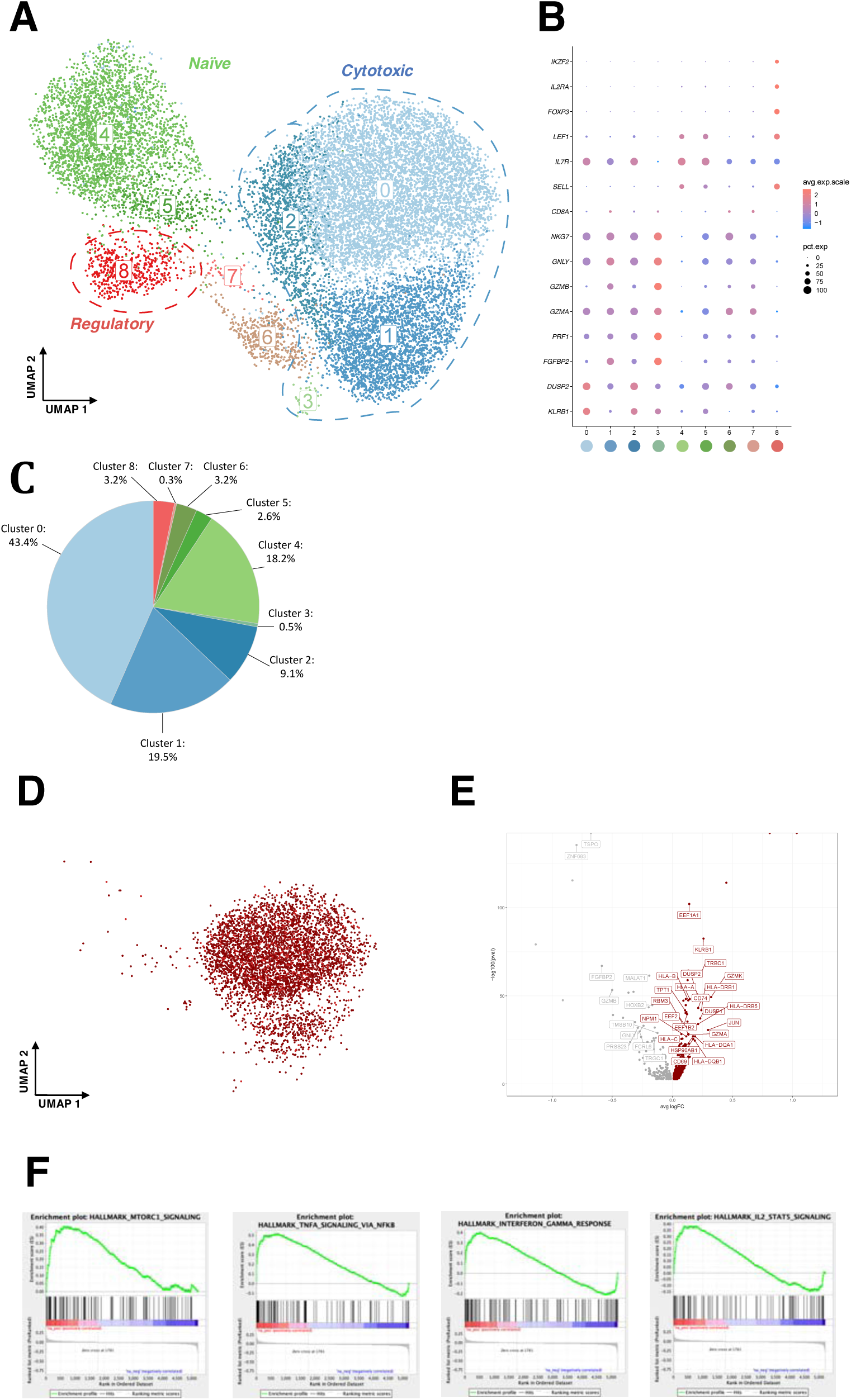
Single-cell RNAseq analysis from the index patient. (A) Two-dimension UMAP-projection of clustered CD4+ cells pooled from two time points from peripheral blood. A total of 15,874 cells are annotated in 9 distinct clusters. B) Gene expression heatmap for the 9 distinct CD4+ clusters, where rows represent canonical marker genes and columns represent different clusters C) Pie chart showing the fractions of cells belonging to different clusters. The fractions are pooled from two time points. D) Graphical visualisation showing the cells taken into differential expression analysis. Red shows the clonally expanded and mutated clonotype, and grey cells represent the cells from other clonotypes with similar phenotype E) Volcano plot showing differentially expressed genes between clonotype of interest and cells from other clonotypes with similar phenotype F) Gene Set Enrichment Analysis results from the differential expression analysis. Shown here are four of eleven HALLMARK-categories enriched (FDR qval < 0.05) to clonotype of interest.

From the TCRab-seq we detected TCRab, TCRa or TCRb from 11,055 cells (71.4%), resulting in 3651 unique T cell clonotypes. The clonotype matching to TCRVB sequencing data (Figure 1C) and harboring the *MTOR*-mutation was also the most expanded fraction, representing 40.1% of the CD4+ T cells. Almost all of the cells from this clonotype (90.1%) belonged to the cytotoxic clusters, and most of the cells were included in the cluster 0 (74.6%) (Figure 4D).

To understand the effect of the *MTOR*-mutation on the T cells, we performed differential expression (DE) analysis between the cytotoxic cells, comparing the cells from clonotype of interest against the other cytotoxic cells in clusters 0, 1, 2 and 3. The analysis found 876 statistically significant DE-genes, of which 694 were upregulated in the clonotype, including cytotoxic genes (e.g. *GZMA*, *GZMAB, GNLY, GZMK*, *NKG7 and PRF1*), and HLA class I and II genes (Figure 4E). Additionally, upregulated eukaryote elongation factors (*eEF*s), such as *EEF1A1, EEF1B2*, and *EEF2*, supported abnormal growth and proliferation of the expanded CD4+ T cells. Furthermore, the expression of *DUSP2* and *KLRB1* genes was highly specific for the mutated clone (Figure 4B). To identify differential pathway regulation in the clonotype, Gene Set Enrichment Analysis (GSEA) was performed, resulting in 11 significantly over-represented and 0 under-represented pathways in the clonotype (Table S11 in the Supplementary Appendix). The upregulated pathways included *MTORC1-*pathway supporting the similar effect of the mutations in patient cells as observed in the *in vitro* cell line models (Figure 4F). Other pathways found in the GSEA analyses included TNF-alpha signaling, IFNg response and IL2-STAT5 signaling.

### Real-time cytotoxicity analysis of CD4+ T cells against primary fibroblasts from the index patient

Real-time electrical impedance measurements monitoring target cell killing have been widely applied to study the cellular cytotoxicity *in vitro*.(*16, 17*) To test the functional effects of the mutated CD4+ T cells and to verify aberrantly upregulated gene expression signatures associated with cytotoxicity, we performed co-culture experiments with CD4+ T cells and primary fibroblasts. Addition of purified CD4+ T cells on the mono-layer of primary fibroblasts from the index patient resulted in decreased electrical impedance implicating cytotoxicity of the CD4+ T cells (Figure 5A and B). In contrast, the CD8+ T cells showed no cytotoxic activity against the fibroblasts, as the impedance curve mirrored the control well without effector cells.

**Figure 5.**
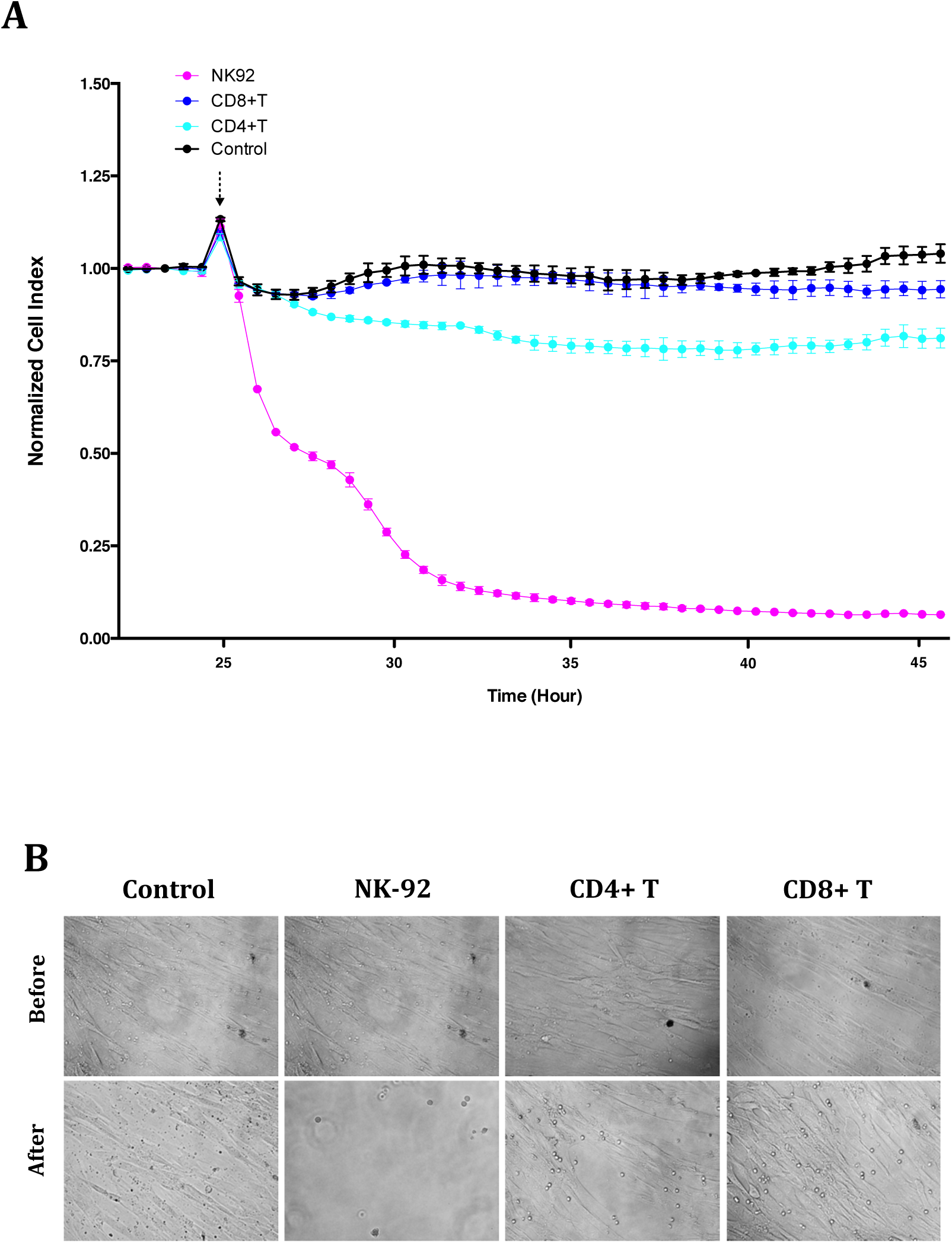
Real-time monitoring of cellular cytotoxicity by electrical impedance measurement. Real-time cell analysing (RTCA) systems, xCELLigence^TM^, was applied to monitor real-time killing effect of primary fibroblasts obtained from the index patient. A) The primary fibroblasts were cultured as monolayers for 24 hours to reach full confluence. Once confluent, the effector cells; NK-92 cell line (positive control; pink line), primary CD4+ T cells (light blue) and primary CD8+ T cells (dark blue) were added to each well and co-cultured (Arrow). The control (black line) shows the impedance of the fibroblasts without any added effectors. The cell impedance was measured every 30 minutes for 48 hours. The measured impedance was expressed as Cell Index with the normalization (n=2). Data is representative of two independent experiments showing similar results. The curve represents the mean Cell Index value from 2 separate wells ± SD. B) Monolayers of the primary fibroblast were visualized before the addition of the effector cells by inverted microscope (Nikon Eclipse TS100)(upper panels). In the end of the experiment, each well was washed with PBS to photograph the live and still attached fibroblast.

### Drug Sensitivity and Resistance Testing (DSRT) in CD4+ T cells from the Index Patient

To determine sensitivity of the mutated cells to targeted therapy, robust *ex vivo* DSRT with 527 drugs in 5 different concentrations (*18*) was performed on freshly isolated CD4+ T cells from the index patient, the donor, and a healthy control (Figure 6). In this screen, the index patient’s CD4+ T cells were less sensitive to mTOR/PI3K inhibitors as compared to the donor’s CD4+ T cells (Figures 6A and B), although constitutive PI3K/AKT/mTOR activity generally predicts rapalog sensitivity. Instead, we observed that heat shock protein 90 (HSP90) inhibitors showed an increased killing effect on the index patient’s CD4+ T cells as compared to CD4+ T cells from the donor and healthy control (Figures 6A, C, and D). Interestingly, scRNA-seq analysis indicated that one of HSP90 family members, HSP90AB1, was significantly upregulated in the expanded CD4+ T cell clonotype, and it was one of the gene expression markers for the clonotype (Figure 4E). With regard to other clinically interesting drug classes, both the donor and the recipient CD4+ T cells were sensitive to HDAC inhibitors, CDK-inhibitors, and proteosome inhibitors. The donor cells were also modestly more sensitive to glucocorticoids (dexamethasone and methylprednisone), but this was not statistically significant. Neither donor nor recipient CD4+ T cells were sensitive to cyclophosphamide, tacrolimus, or methotrexate. However, it should be taken into account that the assay read-out is cell death, and lymphocytes were not activated nor actively proliferating during the experiment.

**Figure 6.**
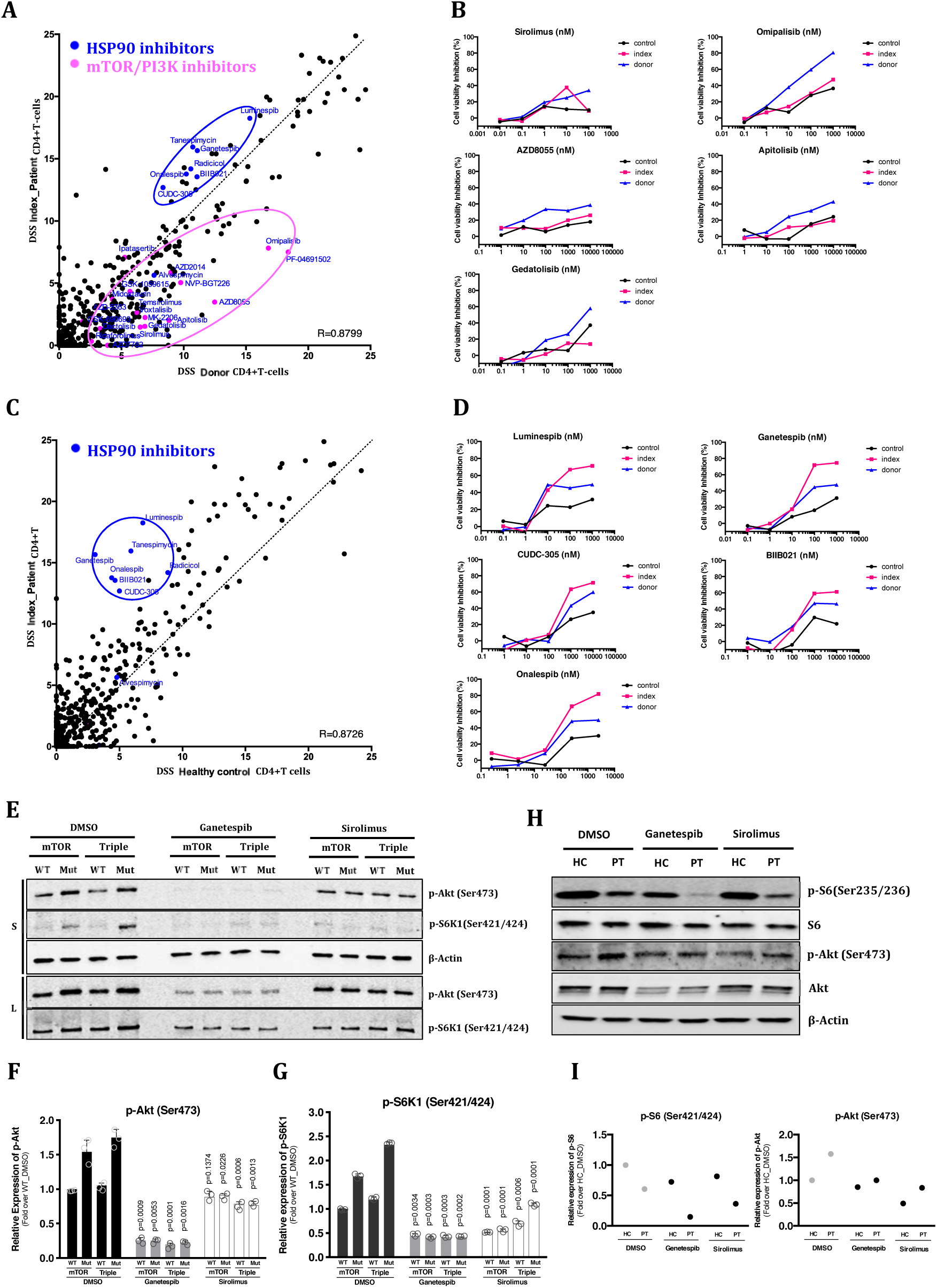
Drug Sensitivity and Resistance testing (DSRT) of CD4+T cells from index patient compared with CD4+T cells from both healthy control and sibling donor. *Ex vivo* DSRT was performed on fresh CD4+T cells from index patient, donor, and healthy control. Correlation of drug sensitivity scores (DSS) indicating cell viability inhibition measured by CellTiter-Glo 2.0 (Promega, USA). DSS is a quantitative measurement of a drug response based on the area under the curve (AUC) with further normalization. Higher DSS denote better killing activity. A) Correlation of DSS scores between index patient and donor CD4+ T cells. B) Individual dose response curves of index patient, donor, and healthy control CD4+ T cells for MTOR inhibitors C) Correlation of DSS scores between index patient and healthy control CD4+ T cells. D) Individual dose response curves of index patient, donor, and healthy control CD4+ T cells for HSP90 inhibitors (E) Stably expressed *MTOR* wildtype (WT), mutant (*P2229R*), Triple (*MTOR, TLR2* and *NFKB2*) wildtype (*WT*) and Triple mutant (*MTOR P2229R*, *TLR2 W559L*, and *NFKB2 P882Q*) (Mut) in HEK293 cells were treated with HSP90 inhibitor (Ganetespib, 100nM) or mTOR inhibitor (Sirolimus, 100nM) for 12 hours. Western blot analysis was performed with the use of anti-pAkt, anti-pS6K1, and anti-β-actin antibodies. Different amount of total protein was loaded in the upper panel (S, 30 ug) and the lower panel (L, 50 ug). Data is representative of three independent experiments. (F and G) Relative expression levels of phospho-Akt (F) and phospho-S6K1 (G) were estimated by measuring each band intensity using ImageJ software (Rasband, W.S., ImageJ, U. S. National Institutes of Health, Bethesda, Maryland, USA, https://imagej.nih.gov/ij/, 1997-2016). Mean ± SEM are shown (n=3 for group). P values are derived from unpaired t-test with Welch’s correction (p value on each bar indicates statistical significance between DMSO and Ganetespib/Sirolimus). (H) Isolated CD4+T cells from healthy control (HC) and index patient (PT) were treated with HSP90 inhibitor (Ganetespib, 100nM) or mTOR inhibitor (Sirolimus, 100nM) for 12 hours. Cells were lysed and proteins (25ug of total protein) were run on the SDS-PAGE gel. Western blot analysis was performed with the use of anti-S6, anti-pS6, anti-Akt, anti-pAkt and anti-β-actin antibodies. I) Relative expression levels of phospho-Akt and phospho-S6 were estimated by measuring band intensity with ImageJ software.

In both WT and mutant HEK293 cells, the HSP90 inhibitor ganetespib reduced AKT phosphorylation on serine 473. As expected, AKT-phosphorylation appeared to be normal following sirolimus treatment as rapalogs only inhibit mTORC1 activity (Figures 6E and F). Treatment with either drug resulted in decreased levels of phosphorylated S6K1 in both mutant and WT cells (Figure 6G). Both drugs also led to a reduced pS6 phosphorylation in CD4+ T-lymphocytes from the index patient and a healthy control (Figures 6H and I), suggesting an inhibitory effect on mTORC1 activity. Likewise, the scRNA-seq data analysis supported that MTORC1 pathway is upregulated in the clonally expanded cytotoxic CD4+ T cells from the index patient (Table S11 in the Supplementary Appendix). AKT was more phosphorylated in the patient’s CD4+ T cells as compared to controls with a slight decrease in both samples upon treatment with ganetespib or sirolimus.

## Discussion

By analyzing blood CD4+ and CD8+ T cells we discovered a recurrent somatic missense mutation in *MTOR* in cGvHD patients. In the index patient, the mutation was limited to the expanded CD4+ T cell clone, it persisted for years and was found in both blood and sclerodermatous skin lesion samples. Paired scRNA- and TCRab-seq verified that the majority of the expanded CD4+ T cells had upregulated expression of genes associated with cytotoxicity and cellular proliferation. Furthermore, mutated CD4+ T cells possessed cytotoxicity against patient’s own primary fibroblasts. No mutations were discovered in the sibling donor samples suggesting the mutations been formed after the allo-HSCT. Functional *in-vitro* studies indicate that the *MTOR* mutation results in a gain-of-function alteration activating both mTORC1 and mTORC2 pathways.

Accumulation of somatic mutations is inherently associated with normal cell division. The role of somatic mutations in cancer is well established, and interestingly, recent reports suggest that somatic mutations may also play a role in the pathogenesis of non-malignant diseases.(*19–23*) We have previously shown that somatic mutations occur in cytotoxic lymphocytes of newly diagnosed rheumatoid arthritis patients(*6*) and in patients with large granular lymphocytic proliferation.(*24*) Interestingly, it was also recently shown, that the disruption of the *TET2* gene by lentiviral vector-mediated insertion of the chimeric antigen receptor (CAR) transgene led to the expansion of single CAR T cell in a patient with chronic lymphocyte leukemia and enhanced therapeutic efficacy.(*25*) In addition to lymphoid cell-related disorders, recent studies have suggested that somatic mutations may play a role in the pathogenesis of other non-cancerous diseases such as *KRAS* mutations in brain arteriovenous malformations leading to activation of the MAPK-ERK pathway.(*23*)

The novel *MTOR P2229R* kinase domain mutation discovered in our cohort has not been previously reported, although already over 750 different *MTOR* mutations exist in the COSMIC database. Previously, mutations in the same domain have been shown to lead to the activation of the mTORC1. The PI3K/AKT/mTOR axis controls important cellular processes and is frequently dysregulated in various cancer types.(*26*) This pathway is also important for the regulation of T cell activation, function, and survival.(*27*) Inhibition of the PI3K/AKT/mTOR axis by targeting the mTORC1 complex with rapamycin has been used to prevent and treat GvHD.(*28*) Based on our data, the *MTOR P2229R* mutation induces the activation of both the mTORC1 and mTORC2 as noted by increased S6K and AKT phosphorylation. Additionally, scRNA-seq analysis strongly supported that *MTORC1* pathway is upregulated in the clonally expanded and cytotoxic CD4+ T cells from the index patient.

In addition to *MTOR*, the mutation in the *NFKB2* gene is also a putative driver due to the importance of this signaling pathway in the immune system. NF-κB has been extensively described as one of the main regulators of the inflammatory response and cancer pathogenesis, offering a promising target in anti-inflammation and -cancer drug development.(*29*) Both p50/RelA-mediated canonical and p52/RelB-mediated non-canonical pathways are involved in NFκB activation. Phosphorylated NFκB2 (p100) is associated with RelB and sequentially ubiquitinated at the C-terminus to form the p52/RelB complex, which is translocating into nucleus and activating downstream target genes.(*30*) Therefore, p52 formation in NFκB2 (p100) processing is a key step for the activation of the non-canonical NFκB pathway. Especially two serine residues in the C-terminal domain of NFκB2 (p100), S866 and S870, are necessary for NFκB2 (p100) processing.(*31*) The index patient harbored *NFkB2* (p100) *P882Q* somatic mutation located in the C-terminus, which has not been reported in the COSMIC before. This mutation leads to an increased p52 formation which potentially induces a hyper-activation of non-canonical NF-κB pathway contributing to chronic inflammation in the index patient.

Unlike the discovered *NFkB2* and *MTOR* mutations, the *TLR2 W558L* somatic mutation was identified in both cGvHD patients and healthy controls. However, VAFs were significantly higher in cGvHD patients as in healthy controls. TLR2 is one of the pathogen-associated molecular pattern recognition receptors and regulators of innate immunity, and it is constitutively expressed on regulatory and memory CD4+ T cells.(*32*) TCR activation by an anti-CD3 antibody has been shown to induce the overexpression of TLR2 on naïve CD4+ T cells followed by activation of the MyD88-dependent *NF-κB* signaling cascade and inflammatory gene expression in peripheral blood.(*33*) Thus, in combination with the *NFkB2* and *MTOR* mutations, the *TLR2 W558L* mutation may further enhance the activation of the *NF-κB* and inflammatory pathways.

Single-cell RNA-seq analysis revealed that the clonally expanded CD4+ T cells had upregulated gene expression signatures associated with cytotoxicity (*GZMA*, *GZMB*, *GNLY* and *NKG7)* and proliferation (*DUSP2*, *KLRB1*, *EEF1A1*, *EEF1B2*, and *EEF2*). Furthermore, real-time impedance analysis indicated cytotoxicity of CD4+ T cells from the index patient, whereas the CD8+ T cells had no effect. Although cytotoxicity has not been considered as a typical CD4+ T cell function (*34*), cytotoxic potential of T helper cells (CD4+CTL) has been recently described including a granzyme-mediated killing capability of target cells during viral infections.(*35, 36*) Epigenetic and molecular mechanisms leading into CD4+CTL differentiation has not yet been clearly described.(*37*) From the proliferation associated genes, DUSP2 (PAC-1) was one of the hallmark genes associated with the mutated clonotype. It is known as one of the MAPK phosphatases, which play an important role in deactivating MAPK. DUSP2 is shown to be up-regulated in T cell activation associated with inflammation.(*38*) Murine knockout phenotype studies for DUSP2 presented reduced cytokine production and protected from inflammatory arthritis. In the murine colitis model, DUSP2 knockout induced Th17 differentiation by directly enhancing the transcriptional activity of STAT3. Therefore, DUSP2 inhibit Th17 lineage of T cell development by attenuated STAT3 activity through dephosphorylation of STAT3 at Tyr705 and Ser727.(*39*) In addition, it has been indicated that DUSP2 inhibit JNK leading to ERK pathway activation.(*40*) Interestingly, TLR2 stimulation also strongly correlate with ERK activation. ERK(1/2) pathway is well known for a positive selection of T cell development and proliferation, and especially CD4+ T cell differentiation depends on ERK signaling.(*41*) Taken together, DUSP2 up-regulation in scRNA seq data support the expansion of CD4+T cells in the index cGVHD patient.

Drug sensitivity testing with patient cells indicated a lower efficacy of mTOR/PI3K inhibitors compared to the donor, suggesting a hyperactivation of mTOR pathway due to the mutation. However, the effect of ongoing immunosuppressive treatment (mycophenolate) cannot be ruled out as it may also affect mTOR signaling.(*42*) Since the mTOR pathway is hyperactivated in many different types of cancer and autoimmune diseases, mTOR inhibitors have been developed and applied to prevent dysregulated mTOR signaling.(*43–46*) Rapamycin and its analogs are highly specific mTOR inhibitors that form a complex with FKBP2, which selectively binds to the FRB domain of mTOR, leading to targeted inhibition of mTORC1-mediated signaling pathways. Therefore, alternative therapeutic applications have been suggested for additionally targeting the dysregulated mTORC2 signaling pathway. Initially, Zheng et al. indicated that the kinase domain of mTOR is a more potent site for mTOR inhibition as it is necessary for both rapamycin-insensitive and rapamycin-sensitive aspects of cell growth and survival.(*47*) ATP-competitive mTOR inhibitors targeting the kinase domain of mTOR have already been developed to inhibit both mTORC1 and mTORC2-mediated signaling processes.(*48, 49*) These inhibitors demonstrated a better clinical efficacy and lower toxicity in anti-tumor therapy as compared with rapalogs.(*7*)

Interestingly, HSP90 inhibitors showed higher efficacy when compared to mTOR inhibitors in the DSRT assay. HSP90 is one of the most abundant and conserved ATP-dependent molecular chaperons, whose expression increases by up to 10-fold under physiologic stress conditions.(*50*) Functionally, HSP90 plays an important role in the refolding of denatured proteins under stress conditions.(*51, 52*) In addition, it activates these proteins including growth-stimulating proteins and kinases.(*50*) mTOR, through its mTORC1 component raptor, directly binds to HSP90 in primary T cells in order to regulate mTOR signaling processes.(*53*) Additionally, AKT physiologically interacts with HSP90 to activate mTOR pathway signaling.(*54*) Notably, HSP90 overexpression has been suggested to be correlated with Akt/mTOR pathway activation in cancer. Furthermore, HSP90 inhibitors have been shown to suppress the Akt/mTOR pathway activity.(*55*) Therefore, HSP90 has become a therapeutic target for several cancer types, and many HSP90 inhibitors are under the evaluation in phase I and II clinical trials for cancer therapy.(*56*) Importantly, HSP90 inhibitors have also been suggested to have protective and therapeutic effects in mouse models of GvHD.(*57*) To our knowledge, these have not yet been used to treat GvHD patients, but based on our results HSP90 inhibitors could serve as a novel therapeutic approach in a subset of cGvHD patients.

In conclusion, novel *MTOR, NFkB2*, and *TLR2* somatic mutations were discovered in an expanded CD4+ T cell clone in a patient with cGvHD. The mutations persisted over time and induced activation of the *NF-kB* and *MTOR* pathways. The *MTOR* mutation was found to be recurrent in other cGvHD patients. Although the *MTOR* mutation frequency was low in the total cGvHD cohort, somatic mutations may also exist in other genes, and similar small subgroups of cGvHD patients can be discovered warranting further investigations. Our findings imply a novel mechanism for the aberrant, persistent T cell activation in cGvHD and pave the way for potential novel individualized therapies.

## MATERIALS AND METHODS

### Study Patients

Samples were collected between 2007-2016 from 135 patients who had developed cGvHD after allo-HSCT (Helsinki University Hospital, Helsinki, Finland, n=8; Turku University Hospital, Turku, Finland, n=37; Hospital de la Princesa, Madrid, Spain, n=19; Hospital Morales Meseguer, Murcia, Spain, n=71). In addition, 38 patients who had not developed cGvHD until the date of sampling, served as a control cohort (Turku University Hospital n=6 and Hospital Morales Meseguer n=32). The blood samples were collected 3 to 102 months (mean 13.5 months) and 2 to 47 months (mean 14.5 months) after allo-HSCT for GvHD and non-cGvHD patients, respectively. Clinical characteristics of the patients are summarized in Supplemental Table S1. All patients provided written informed consents. Additionally, buffy coat samples from 54 healthy blood donors were obtained from the Finnish Red Cross Blood Service. A peripheral blood sample from the index patient’s sibling donor was also obtained.

The study was performed in compliance with the principles of Helsinki declaration, and was approved by the ethics committees in the Helsinki University Hospital (Helsinki, Finland), Turku University Hospital (Turku, Finland), Hospital de la Princesa (Madrid, Spain) and Hospital Morales Meseguer (Murcia, Spain).

### Reagents

Primary antibodies against NFkB2 (Cat#: 4882S, Lot: 4), ribosomal protein S6 (Clone: 54D2, Cat#: 2317S, Lot#: 4), phospho-S6 ribosomal protein (Ser235/236) (Clone: D57.2.2E, Cat#: 4858T, Lot#: 16), Akt (Clone: C67E7, Cat#: 4691T, Lot#: 20), phospho-Akt (Ser473) (Cat#: 9271T, Lot#: 14), p70S6 kinase (S6K) (Clone: 49D7, Cat#: 2708T, Lot#: 7), and phospho-p70S6 (S6K) kinase (Thr421/Ser424) (Cat#: 9204S, Lot#: 11) were purchased from Cell Signaling Technology, and antibody against beta-actin (Clone: AC15, Cat#: ab6276, Lot#: GR66278-11) was purchased from Abcam. Secondary antibodies, IRDye 700 conjugated anti-mouse and IRDye 800 conjugated anti-rabbit, were purchased from LI-COR Biosciences.

### Cell Lines

Human embryonic kidney HEK293 cells (Cat#: CRL-1573, ATCC) and HEK293FT (Cat#:R70007, Thermo Fisher Scientific) were maintained in high-glucose Dulbecco Modified Eagle medium (Lonza) containing 10% FBS (Gibco), 1% penicillin-streptomycin (Invitrogen), and L-glutamine (Lonza) in a 37 °C humidified incubator with 5% CO2. Cell line authentication was performed with GenePrint10 System (Promega). HEK293 cell line was authenticated *via* Promega GenePrint10 System. The result was compared ATCC STR, JCRB STR, ICLC STR database, and DSMZ online STR database. The identity estimates are calculated according to the allele information found in these databases. HEK293FT cell line was derived from Thermo Fisher Scientific with an authentication and only less than 10 passages were used. Mycoplasma test was performed using MycoAlert Mycoplasma Detection Kit (LONZA, Cat#: LT07-318).

### Sample Preparation and DNA Extraction

Mononuclear cells (MNCs) were separated from whole blood using Ficoll-Paque^TM^ PLUS (GE Healthcare). The separated MNCs were then labeled with either CD4+ or CD8+ magnetic beads (Miltenyi Biotec) and sorted by AutoMACs® cell sorter (Miltenyi Biotec) according to the manufacturer’s protocol. The purity of sorted fractions was evaluated by flow cytometry and confirmed to be >98% (FACsVerse, BD Biosciences). Alternatively, separated MNCs were sorted using FACsAria II (BD Biosciences). Genomic DNA was isolated from fresh or frozen sorted MNCs or from whole blood samples using the Genomic DNA NucleoSpin Tissue kit (Macherey-Nagel). DNA concentration and purity were measured with Qubit2.0 Fluorometer (Invitrogen) or Nanodrop (Thermo Fisher Scientific).

### Flow Cytometry Analysis and Flow-assisted Cell Sorting

For phenotyping of the memory T cell subsets, peripheral blood mononuclear cells (PBMCs) were immunostained with the antibody panel including anti-CD3 PeCy7 (Clone: SK7, Cat#: 557851, Lot#: 8037645, BD Biosciences), -CD4 PerCP (Clone: SK3, Cat#: 345770, Lot#: 6281605, BD Biosciences), -CD8 PerCP (Clone: SK1, Cat#: 345774, Lot#: 82152, BD Biosciences) -CD45RA Alexa700 (Clone: HI100, Cat#: 560673, Lot#: 7180940, BD Biosciences), and -CCR7 PE (Clone: 150503, Cat#: FAB197P, Lot#: LEU1618031, R&D System). Stained samples were analyzed with FACSVerse (BD Biosciences) and FlowJo software (Version 10.4.2). For CD4+ T cell TCR Vβ20+ clone sorting, PBMCs were immunostained with anti-CD3 APC (Clone: SK7, Cat#: 345767, Lot#: 7236657, BD Biosciences), -CD4 PerCP (Clone: SK3, Cat#: 345770, Lot#: 6281605, BD Biosciences), -CD8 PE-Cy7 (Clone: SK1, Cat#: 335822, Lot#: 8272690, BD Biosciences) and – Vβ20 (IOTest® Beta Mark TCR Vbeta Repertoire Kit, Cat#: IM3497, Lot#: 66, Beckman Coulter). Stained cells were physically isolated by FACs AriaIII (BD Biosciences) and analyzed with FlowJo software. Purity of sorted cells was more than 99% and verified with the same system. (Supplementary Figure S2C)

### TCR V**β** Analysis

TCR Vβ families were analyzed from peripheral whole-blood samples by flow cytometry based antibody staining using IOTest® Beta Mark TCR Vβ Repertoire Kit (Cat#: IM3497, Lot#: 66, Beckman Coulter). Briefly, CD4+ and CD8+ T cells in whole blood samples were stained with the panel of TCR Vβ antibodies recognizing 24 members of TCR β chain, which covers about 70% of the normal human TCR Vβ repertoire. Stained cells were further analyzed using FACSVerse (BD Biosciences).

### TCR CDR3 Deep Sequencing

Isolated genomic DNAs was used for TCRB deep sequencing. Sequencing and data analysis was conducted with ImmunoSEQ assay as previously described (Adaptive Biotechnologies, Seattle, WA).(*58*)

### Immunopanel Sequencing

A customized NGS panel including exonic areas of 986 genes related to immunity and cancer was used to screen for somatic mutations.(*6*) Genes included in the panel are provided in the Supplemental Table S2. Sequencing was done from both sorted CD4+ and CD8+ T cells. Bioinformatic analysis to identify and annotate somatic variants was performed as previously described.(*6, 24*)

### Validation of the Somatic MTOR Mutation by Capillary Sequencing Analysis

A specific primer set was designed using the Primer-Blast search (National Center for Biotechnology Information: http://blast.ncbi.nlm.nih.gov/) to validate the somatic MTOR mutation (Supplementary Table S3). Polymerase chain reaction (PCR) products were purified with the ExoSAP-IT (Affymetrix) followed by sequencing on DNA sequencer (Applied Biosystems). Sequences were analyzed using 4Peaks version 1.7.1.

### Amplicon Sequencing of MTOR, NFkB2 and TLR2

Targeted amplicon sequencing was performed with an in-house developed deep amplicon sequencing panel using the Illumina Miseq platform (Supplementary Table S4). The coverage was over 100,000 X, and a variant was called if variant base frequency was 0.5% of all reads covering a given a position. All variants with the base quality frequency ratio (ratio of number of variant calls/ number of all bases and quality sum of variant calls / quality sum of all bases at the position) ≥ 0.9 were considered as true somatic variants. A detailed sequencing protocol and the bioinformatics pipeline used for data analysis are described in previous reports.(*6, 24*)

### scRNA-seq and TCRab-seq analysis

CD4+ T cells from two time points of the index patient were enriched using CD4 microbeads (Miltenyi Biotec). Single cells were partitioned using a Chromium Controller (10x Genomics) and scRNA-seq and TCRab-libraries were prepared using Chromium Single Cell 5’ Library & Gel Bead Kit (10x Genomics), as per manufacturer’s instructions (CG000086 Rev D). In brief, 17,000 cells from each sample, suspended in 0.04% BSA in PBS were loaded on the Chromium Single Cell A Chip. During the run, single-cell barcoded cDNA is generated in nanodroplet partitions. The droplets are subsequently reversed and the remaining steps are performed in bulk. Full length cDNA was amplified using 14 cycles of PCR (Veriti, Applied Biosystems). TCR cDNA was further amplified in a hemi-nested PCR reaction using Chromium Single Cell Human T Cell V(D)J Enrichment Kit (10x Genomics). Finally, the total cDNA and the TCR-enriched cDNA was subjected to fragmentation, end repair and A-tailing, adaptor ligation, and sample index PCR (14 and 9 cycles, respectively). The gene expression libraries were sequenced using an Illumina NovaSeq, S1 flowcell with the following read length configuration: Read1=26, i7=8, i5=0, Read2=91. The TCR-enriched libraries were sequenced using an Illumina HiSeq2500 in Rapid Run mode with the following read length configuration: Read1=150, i7=8, i5=0, Read2=150. The raw data was processed using Cell Ranger 2.1.1. with GRCh38 as the reference genome.

During secondary analysis, cells with fewer than 200 or more than 4000 genes, or more than 15% of the counts from mitochondrially-encoded transcripts were excluded from the analysis. The remaining data was log-normalized and scaled. To reduce the dimensionality of the data, we determined the highly variable genes as the genes with the highest variance-mean ratio. Genes that had mean expression between 0.0125 and 3 on a log-transformed count scale and genes above 0.5 log(variance/mean) were counted as highly variable, resulting in 764 genes. The T-cell receptor V-genes, mitochondrial genes and ribosomal genes (n = 113) were excluded from the results, resulting in a final list of 651 highly variable genes.

Clusters were identified using the graph-based community identification algorithm as implemented in the Seurat-package(*28*). Prior to calculating cell-cell distances, PCA was performed on the 651 highly variable genes on all the QC-positive cells, and the top 50 principal components were kept. To prevent overclustering, the optimal number of clusters was determined by increasing the resolution hyperparameter as a function of number of clusters until the first saturation plateau was achieved. The robustness of these clusters was assessed by subsampling cells and doing the analysis iteratively and visually inspecting the results of embedding and differentially expressed genes between the formed clusters. Differential expression analysis was performed based on the t-test, as suggested by Robinson et al(*59*). Clusters were annotated using canonical cell type markers as well as the differentially expressed genes.

Gene Set Enrichment Analysis (GSEA) (software.broadinstitute.org/gsea/index.jsp) between the clonotype and other cytotoxic cells was performed on genes that were detected at least in 0.1% of the cytotoxic cells and had at least log fold-change of 0.01 between the clonotype and other cytotoxic CD4+ T cells. The gene list was ordered based on the fold-change. Overlap with HALLMARK-category was assessed and the False Discovery Rate (FDR) calculated while the number of permutations was 1000.

Clonotypes were identified based on the available information, and both total nucleotide level TCRa and TCRb were used if found. Cells for which more than two recombinants were identified were excluded from further analysis.

From the TCRab-seq we detected TCRab, TCRa or TCRb from 11,055 cells (71.38%), resulting in 3651 different T cell clonotypes. The clonotype harboring the *MTOR*-mutation was the most expanded, compromising of 2366 cells (TRA:CLVGDIGNQGGKLIF; TRB:CAWSTGQANNSPLHF). However, we noticed that the second most (TRB:CAWSTGQANNSPLHF, 1545 cells) and third most expanded (TRA:CLVGDIGNQGGKLIF, 598 cells) clonotypes had only one chain and as they matched to the most expanded clonotype, we treated this as error coming from uncomplete sequencing and pooled the three most expanded clonotypes into one.

### Analysis of cellular cytotoxicity

Primary fibroblasts from the index patient were cultured based on previous method.(*60*) Briefly, skin biopsy from the index patient were dissected in small pieces (approx. 2mm × 2mm) and transferred into 6-well plate in 500 µl of complete growth medium containing 20% FBS. 2-300µl of growth medium was added for every 2 days to replace evaporated media. After one week, increase amount of media to 2 ml and change the media every 3 days. Once cells were confluent in each well, cells were trypsinized and passaged.

To measure cellular cytotoxicity of CD4+ and CD8+ T cells from the index patient, the proliferation of the fibroblast established from the index patient was monitored with xCELLigence^TM^ real-time cell analyzer (RTCA) (ACEA Biosciences, CA, USA) according to the manufacture’s instruction. xCELLigence^TM^ RTCA biosensor measures cellular adhesion through electrical impedance, which is converted to Cell Index (arbitrary units). Briefly, the E-Plate 16 VIEW (ACEA Biosciences, CA, USA) was equilibrated with the 100 µl of culture media at room temperature. 100 µl of the cell suspension (8 × 10^3^ cells/well) in duplicate was transferred to the plate followed by incubation at room temperature for 30 min to allow the cells to settle at the bottom of the wells. The xCELLigenceTM monitored the cells every 30 min for 200 repetitions. When the cell index [(impedance at time point n – impedance in the absence of cells)/nominal impedance value] were reached a plateau, CD4+ T cells, CD8+ T cells, and NK92 as an cellular cytotoxicity inducer (6.4 × 10^4^ cells as a ratio of the fibroblast to the inducer is 1:8) for the fibroblast were added on the plate. CD4+ T cells and CD8+ T cells were separated from MNCs with CD4+ or CD8+ magnetic beads (Miltenyi Biotec) and sorted by AutoMACs® cell sorter (Miltenyi Biotec) according to the manufacturer’s protocol. The real-time impedance trace for the fibroblasts exposed to CD4+ T cells, CD8+ T cells, and NK-92 were monitored for 48 h.

### Multiplexed Immunohistochemistry (mIHC)

Tissue blocks were cut in 3.5 µm sections. Slides were deparaffinized in xylene and rehydrated in graded ethanol series and H2O. Heat-induced epitope retrieval (HIER) was carried out in 10 mM Tris-HCl – 1 mM EDTA buffer in +99°C for 20 min (PT Module, Thermo Fisher Scientific). Peroxide activity was blocked in 0.9% H2O2 solution for 15 min, and protein block performed with 10% normal goat serum (TBS-NGS) for 15 min. Anti-CD3 (Clone: EP449E, Cat#: ab52959, Lot#: GR140731, Abcam) primary antibody diluted 1:500 in protein blocking solution and secondary anti-rabbit horseradish peroxidase-conjugated (HRP) antibodies (Immunologic) diluted 1:1 in washing buffer were applied for 1h45min and 45 min, respectively. Tyramide signal amplification (TSA) 488 (PerkinElmer) was applied on the slides for 10 min. Thereafter, HIER, peroxide and protein block were repeated, followed by application of anti-CD8 (1:500, Clone: C8/144B, Cat#: BSB 5174, Lot#: 5174JDL05, BioSB) primary antibody, HRP-conjugated secondary antibody diluted 1:3 with washing buffer and TSA 555 (PerkinElmer). HIER, peroxide block and protein block were repeated. Then, the slides were incubated with CD4 primary antibodies (1:25, Clone: EPR6885, Cat#: ab133616, Lot#: GR218457, Abcam) overnight in +4°C. Next, AlexaFluor647 fluorochrome-conjugated secondary antibody (Thermo Fisher Scientific) diluted in 1:150 and Dapi (Roche) counterstain diluted 1:250 in washing buffer were applied for 45 min. ProLong Gold mountant (Thermo Fisher Scientific) and a coverslip were applied on the slides. After peroxide block, antibody incubations and fluorochrome reaction, slides were washed three times with 0.1% Tween-20 (Thermo Fisher Scientific) diluted in 10 mM Tris-HCL buffered saline pH 7.4 (TBS). Fluorescent images were acquired with the AxioImager.Z2 (Zeiss) microscope equipped with a Zeiss Plan-Apochromat 20x objective.

### Site-Directed Mutagenesis

Site-directed mutagenesis was conducted using GENEART® Site-Directed mutagenesis system according to the manufacturer’s instruction (Invitrogen) with NFkB2 (GeneCopoeia, Cat.No. EX-Z4293-Lv154), TLR2 (GeneCopoeia, Cat.No. EX-Q0161-Lv122, GeneCopoeia), and mTOR (Addgene, Cat.No.26603) expression vector. The primer sequences used for the site-directed mutagenesis are in the Supplemental Table S5.

### Establishing Stable Cell Lines

HEK293 cells were transfected using FuGENE HD transfection reagent (Promega) with either a wildtype or P2229R *MTOR* expression vector (ratio of reagent to DNA is 3:1) following the manufacturer’s instruction. Neomycin resistant clones were selected after the cells were cultured with G418 (500 µg/mL) for 3 weeks. The lentiviruses were produced by co-transfection of HEK293FT cells with *NFkB2* (wildtype or P882Q mutant) or *TLR2* (wildtype or W558L mutant) lentiviral expression vectors, and psPAX2 lentiviral packaging plasmid (Addgene) and pCMV-VSV-G envelope plasmid (Addgene) using Lipofectamine® 2000 (Thermo Fisher Scientific). Antibiotic free DMEM containing 10% FBS was used as a culturing medium and Opti-MEM I Reduced Serum Medium (Thermo Fisher Scientific) supplemented with 5% FBS and 1 mM Sodium pyruvate was used as a lentivirus packaging medium. 6 hours post-transfection, medium was removed and replaced with DMEM. After 48 hours, the supernatants were centrifuged at 300 g for 5 min to remove cell debris and filtered with a 0.45 µm polyethersulfone membrane filter. Ultracentrifugation to concentrate the virus was performed for 2 hours at 12,000 g and 4°C using Beckman SW28 rotor. Lentivirus titers were measured by p24 specific enzyme-linked immunosorbent assay.

### Establishment of Triple Mutant Stable Cell Lines

HEK293 cells stably expressing exogenous *MTOR* (wildtype or P2229R mutant) were transduced with NFkB2 (wild type or P882Q mutant) expressing lentiviruses. Infections were performed in the presence of 8 µg /mL of polybrene under centrifugation (500 g, 37°C) for 2 hours. *MTOR-NFkB2* transduced cells (expressing Cyan Fluorescent Protein) were selected by using FACsAriaIII (BD Biosciences). Cells expressing exogenous *MTOR-NFkB2* (wildtype or mutant) were infected with *TLR2* (wildtype or W558L mutant) expressing lentiviruses as described above and selected using puromycin (3 µg/mL).

### Western Blot Analysis

After removing serum containing medium, HEK293 cells were washed twice with ice-cold PBS followed by serum starvation for 12 hours. Cells were then harvested and further lysed in ice-cold RIPA buffer with 1X protease and phosphatase inhibitor cocktail (Thermo Fisher Scientific). To remove cell debris, centrifugation was carried out for 10 min at 4 °C, 12,000 g. Total protein concentration was measured with the Qubit protein assay (Thermo Fisher Scientific) and 5 µg of protein per sample was prepared in Laemmil buffer (Bio-Rad Laboratories) to load on a SDS-PAGE gel (Bio-Rad Laboratories). After running the sample in the SDS-PAGE gel, the proteins were transferred into a nitrocellulose membrane (Merk Millipore) followed by blocking the membrane with Odyssey blocking buffer (LI-COR Biosciences) for 1 hour. Primary antibodies (1:1000 dilution) were incubated overnight at 4°C in PBS with 0.1% Tween 20 containing 5% milk, and subsequently secondary antibodies (1:15,000 dilution) in PBS with 0.1% Tween 20 containing 5% milk were incubated for 1 hour at room temperature. The proteins were visualized using Odyssey Imaging Systems (LI-COR Biosciences).

### Drug Sensitivity and Resistance Testing (DSRT)

*Ex-vivo* DSRT was performed on freshly isolated CD4+ T cells with a total of 527 drugs in 5 concentrations covering a 10,000-fold concentration range including conventional chemotherapeutics and a broad range of targeted oncology compounds.(*47*) To dissolve the drug compounds, 5 µl of medium was dispensed into each well of 384 well plates including five different concentrations of each drug. 20 µl of cell suspension (CD4+ T cells from healthy control, donor and index patient: 2,000 cells per well) was transferred to every well using MultiFlo FX dispenser (BioTek). After incubation (5% CO2 at 37°C) for 72 hours, the cell viability was evaluated by CellTiter-Glo Assay solution (Promega). The drug sensitivity score (DSS) was calculated to evaluate quantitative drug profiles based on the measured dose-response curve.(*61*)

### Reverse Transcriptase-quantitative Polymerase Chain Reaction (RT-qPCR)

Total RNA was extracted using the RNeasy Mini kit (Qiagen) followed by cDNA synthesis using QuantiNova Reverse Transcription kit (Qiagen) according to the manufacturer’s protocol. The cDNA was applied in SYBR Green RT-PCR master mix (Applied Biosystems) and oligonucleotide primers (Supplementary Table S6). All RT-qPCR reactions were performed in 384-microwell plates (Applied Biosystem) using a QuantStudio 6 Flex Real-Time PCR system (Applied Biosystems). The relative quantitation of gene expression was analyzed using comparative cycle threshold (ΔΔCT) method, and beta actin (ACTB) was used as an endogenous control to normalize gene expression level.

### Data availability

Patient whole-exome and RNA-sequencing raw data related to table 1 and figure 4 are available from the corresponding author upon request owing to regulations pertaining to the authors ethics permit and deposition of these data in public repositories.

### Statistical Analysis

Unpaired two-sided t-tests were performed using GraphPad Prism 6 for Mac OS X, version 6.0. In all analyses, *P*-value < 0.05 was considered as statistically significant.

## List of Supplementary Materials

Supplementary results: Clinical Characteristics of the Index cGvHD Patient

Supplementary Figure S1. Medical history of the index patient

Supplementary Figure S2. Flow cytometry analysis of the index patient

Supplementary Figure S3. Variant allele frequencies of TLR2 mutations in cGVHD patients’ and healthy controls’ CD4+ and CD8+ T cells

Supplementary Figure S4. Functional analysis of wild type and TLR2 mutants in HEK293 cell line

Supplementary Figure S5. Gene expression fold change within CD4+ cell clusters between 2015 and 2017.

Supplementary Table S1. Summary of study cohorts

Supplementary Table S2. Gene list in the immunogene panel sequencing.

Supplementary Table S3. Primer sets of mTOR, NFkB2 and TLR2 amplicon sequencing

Supplementary Table S4. Primer set for mTOR, NFkB2 and TLR2 capillary sequencing

Supplementary Table S5. List of mTOR, NFkB2 and TLR2 mutagenesis primers

Supplementary Table S6. Primer list of RT-qPCR

Supplementary Table S7. Immunogene panel sequencing result for the index patient

Supplementary Table S8. Somatic NFKB2, TLR2 and MTOR mutations in different cellular fractions from the index patient validated by amplicon sequencing

Supplementary Table S9. Somatic TLR2 mutation in CD4+ T-cells validated by amplicon sequencing

Supplementary Table S10. Somatic TLR2 mutation in CD8+ T-cells validated by amplicon sequencing

Supplementary Table S11. Gene set enrichment analysis (GSEA)

## Acknowledgements

**Funding:** This work was supported by the European Research Council (M-IMM project), Academy of Finland, Finnish special governmental subsidy for health sciences, research and training, Sigrid Juselius Foundation, Instrumentarium Science foundation, Helsinki Institute of Life Sciences Fellow funding, and Finnish Cancer Institute. TL was supported by the Academy of Finland (Decisions 311081 and 314557). This study was supported by Finnish Functional Genomics Centre, University of Turku, Åbo Akademi University and Biocenter Finland.

**Author Contributions:** G.P., D.K., S.L., R.K., M.K. and S.M. designed the study and experiments. A.M.H., C.M-C., L.C., V.G.G.S., T.H.C-L., A.K., A.J., U.S., and M.I-R. contributed and prepared biological samples and clinical data. G.P., D.K., S.L., R.K., O.B., J.H., and T.L. performed experiments and analyzed the data. P.E. and S.H. designed, supervised and performed sequencing assays. S.E. performed bioinformatic analyses. S.M. conceived and designed the study, directed and supervised research. G.P, M.K., and S.M. wrote the manuscript. All authors read and approved the final manuscript.

**Competing Interests statement:** S.M. has received honoraria and research funding from Novartis, Pfizer and Bristol-Myers Squibb (not related to this study). The remaining authors declare no competing interests.

**Data and materials availability:** Patient whole-exome sequencing and RNA-seq data are available from the corresponding author upon suitable request. All other data associated with this study are available in the main text or the supplementary materials.

