## supplementary data for "Somatic Mutations in Clonally Expanded T-lymphocytes in Patients with Chronic Graft-Versus-Host Disease" (from another version of this preprint: version 1, 2019-08-28)

**Supplementary Appendix**

**Contents:**

**Supplementary results** **page 2**
**Supplementary Figure S1** **page 3**
**Supplementary Figure S2** **page 4, 5**
**Supplementary Figure S3** **page 6**
**Supplementary Figure S4** **page 7**
**Supplementary Figure S5** **page**
**Supplementary Table S1** **page**
**Supplementary Table S2** **page**
**Supplementary Table S3** **page**
**Supplementary Table S4** **page**
**Supplementary Table S5** **page**
**Supplementary Table S6** **page**
**Supplementary Table S7** **page**
**Supplementary Table S8** **page**
**Supplementary Table S9** **page**
**Supplementary Table S10** **page**
**Supplementary Table S11** **page**

### Supplementary results

#### Clinical Characteristics of the Index cGvHD Patient

The index patient was a 56-year-old male, who was diagnosed with chronic phase chronic myeloid leukemia (CML) in the fall 1999. The clinical status and treatment history are described in detail in Supplemental Figure 1. After short leukocyte reduction with hydroxyurea and the patient received allo-HSCT from his HLA-matched sibling donor after myeloablative conditioning. Double immunosuppression with cyclosporine and methylprednisone was started at the time of the allo-HSCT, and methotrexate was also given on day +1, +3 and +6 after the transplantation. Within the first 100 days the patient had a human herpesvirus 6 (HHV6) infection and the first cytomegalovirus (CMV) reactivation. At the end of that period, the first symptoms of GvHD emerged in the form of elevated transaminases. Since the beginning of 2001, the patient suffered from cGvHD affecting his liver, eyes, nails and skin including sclerodermatous skin lesions that presented with a varying degree of severity over time. Mycophenolate was added to the immunosuppressive regimen in the spring 2001. In the course of the cGvHD, the patient was treated with photopheresis in 2002 as well as with a low-dose irradiation of his lymph nodes in 2003. The immunosuppression was continuously adjusted according to the clinical presentation of the cGvHD. In recent years, the patient has mostly received mycophenolate and varying supplementation with methylprednisone and cyclosporine depending on the manifestation of sclerodermatous skin lesions. The patient haven’t had a relapse of CML nor received any donor lymphocyte infusions. He was sampled (blood) for the first time in 2013 and in the disease course thereafter (Supplementary Figure S1).

### Supplementary Figure S1. Medical history of the index patient

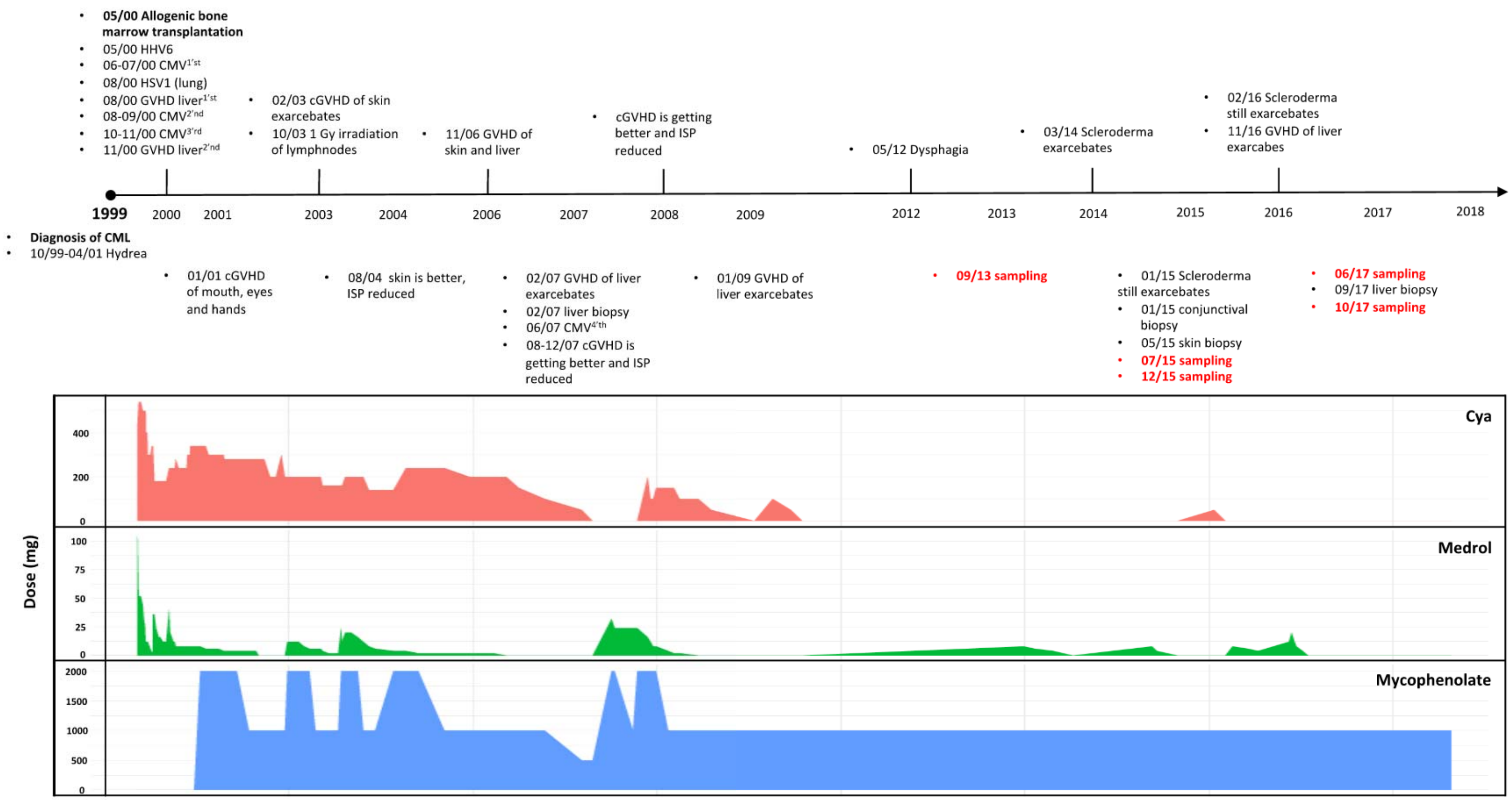

### Supplementary Figure S2.

#### A Lymphocyte population in peripheral blood

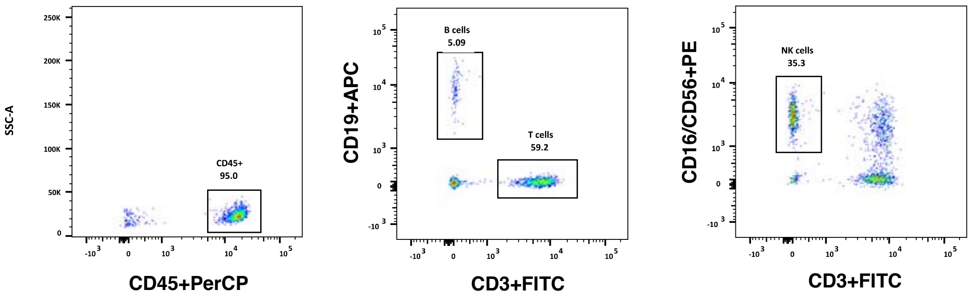


#### B Phenotype of donor and index patient's T cells

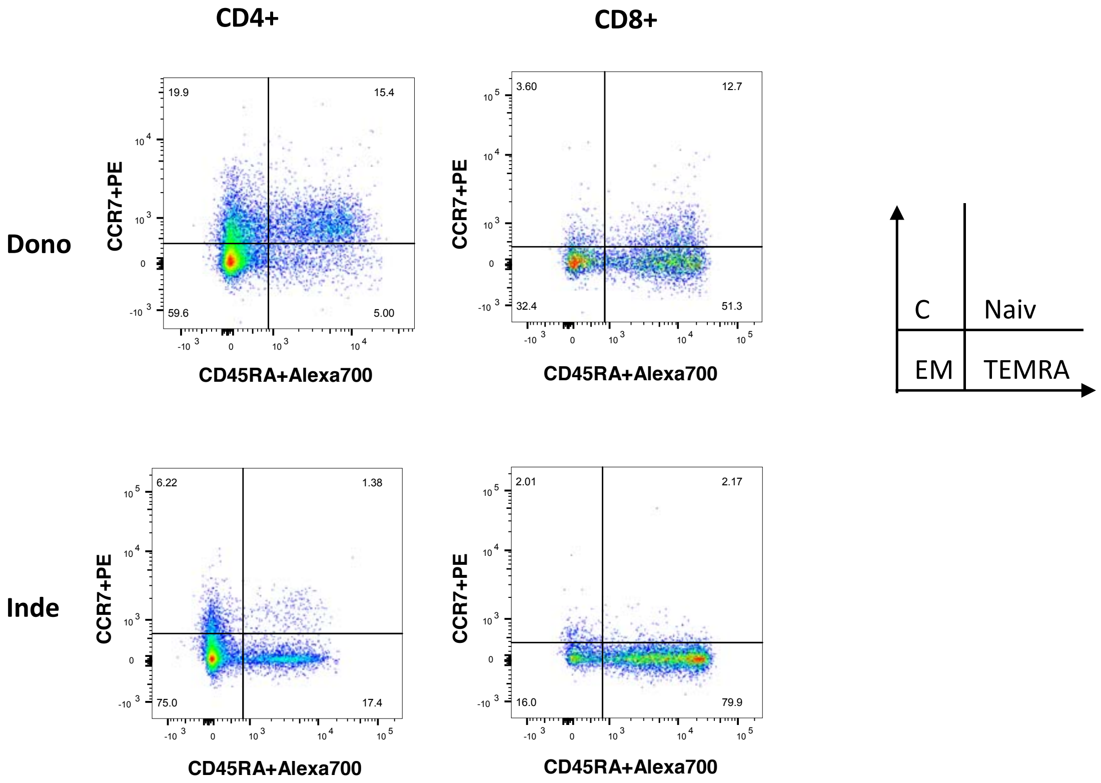

### C Gating strategy for flow-assisted cell sorting

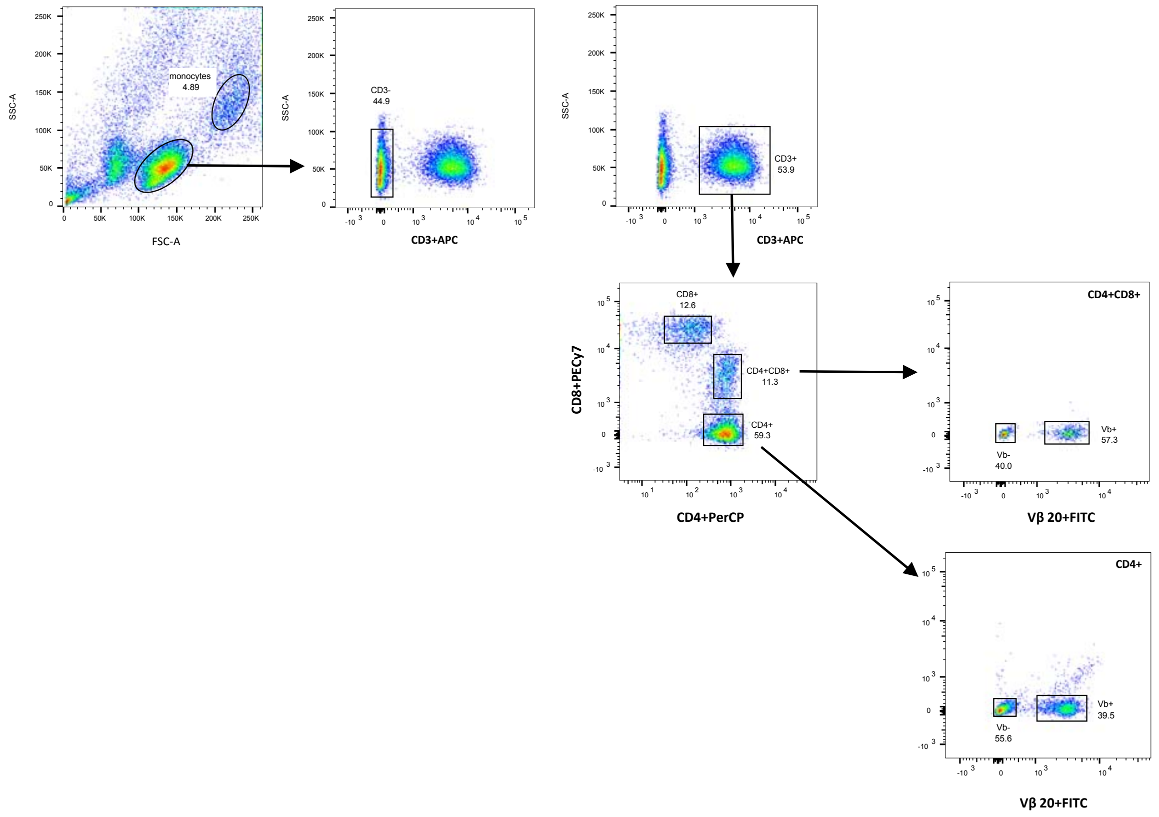


**Supplementary Figure S2. Flow cytometry analysis of the index patient.**
(A) Basic lymphocyte immunophenotyping was performed to identify T (CD3+), natural killer (NK) (CD3-CD16+CD56+), and B (CD3-CD19+) cells by flow cytometry.
(B) Multicolor-flow cytometry was applied to identify the detailed immunophenotype of donor and index patient's memory T cell subtypes. Central memory (CM), naive, effector memory (EM) and terminal effector-memory (TEMRA) cells.
(C) Gating strategy for flow based cell sorting. Monocytes were gated based on morphological features using SSC-A and FSC-A plots. CD3- and CD3+ cells were sorted from lymphocytes based on antibody staining, and CD3+ cells were separated to CD4+, CD4+CD8+ and CD8+ fractions. Finally, TCR Vβ20+ and Vβ20- cells were sorted.

**Supplementary Figure S3.**

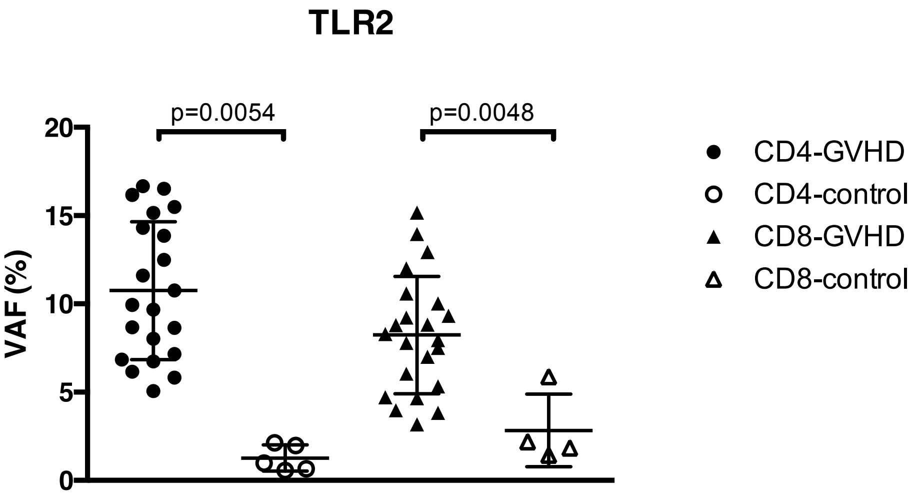


**Supplementary Figure S3. Variant allele frequencies of TLR2 mutations in cGVHD patients' and healthy controls' CD4+ and CD8+ T cells**

TLR2 mutation analyzed by amplicon sequencing in cGVHD patients (CD4+ T, n=21 and CD8+ T, n=22) and healthy controls (CD4+ T, n=5 and CD8+ T, n=4). Mononuclear cells (MNCs) were separated from whole blood with Ficoll-Paque PLUS (GE Healthcare, UK) followed by magnetic beads sorting of both CD4 and CD8+ T cells. The purity of sorted fractions was evaluated by flow cytometry (FACsVerse) and confirmed to be >98%.

The analysis indicated 10-fold higher variant allele frequency in cGVHD patients compared to healthy controls. Unpaired t-test was performed with Welchi's correction to calculate p value using Graphpad Software (San Diego, USA). Error bar present Mean ± SD.

**Supplementary Figure S4.**

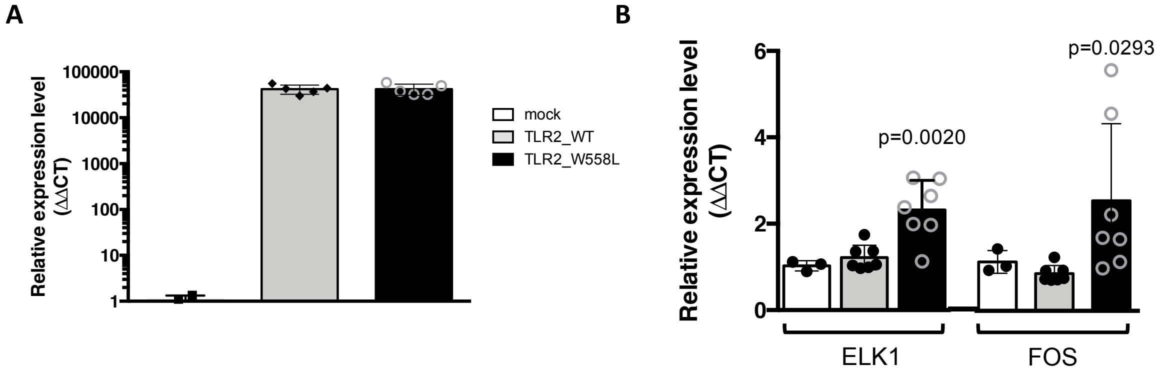


**Supplementary Figure S4. Functional analysis of wild type and TLR2 mutants in HEK293 cell line**

To identify the effect of TLR2 mutant on the downstream gene expression, RT-qPCR was performed for (A) TLR2 and (B) ELK1 and FOS1 genes using stably expressed TLR2_WT (n=7) and TLR2_W558L (n=7) in HEK293 cells.

TLR2_W558L increased expression level of downstream target genes, ELK1 and FOS1. Unpaired t-test was performed with Welchi's correction to calculate p value (TLR2_WT vs TLR2_W558L) using Graphpad Software (San Diego, USA). Error bar present Mean ± SD.

**Supplementary Figure S5.**

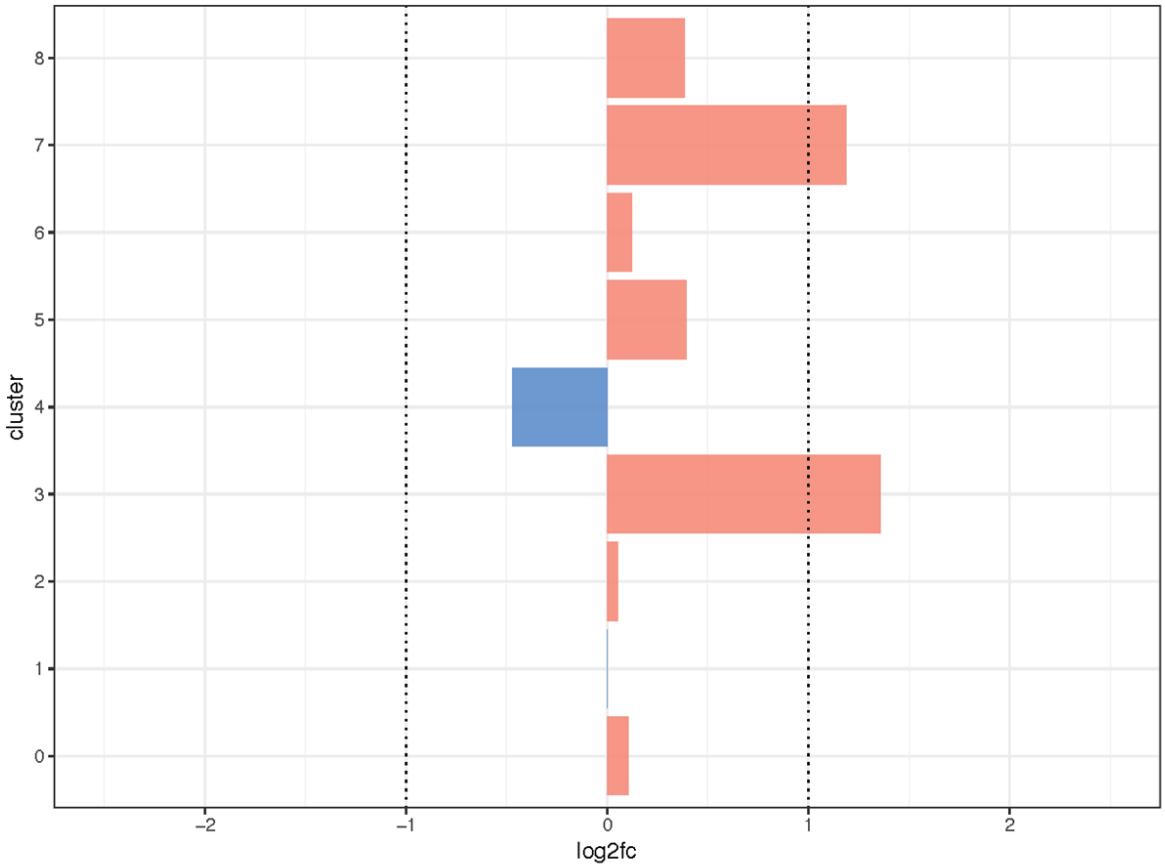


**Supplementary Figure S5. Gene expression fold change within CD4+ T cell clusters between 2015 and 2017**

**Supplementary Table S1. Summary of Study Cohorts**

**A. Patients with cGvHD**

| Patient characteristics | N | % |
|---|---|---|
| **Total** | 135 | |
| **Age at sampling (mean, range)** | 48 | (16-70) |
| **Patient sex** | | |
| Female | 58 | 43,0 |
| Male | 77 | 57,0 |
| **Donor sex*** | | |
| Female | 53 | 39,3 |
| Male | 81 | 60,0 |
| **Sex mismatch*** | | |
| No | 73 | 54,1 |
| Yes | 61 | 45,2 |
| **Diagnosis** | | |
| AML, MDS | 63 | 46,7 |
| ALL | 17 | 12,6 |
| NHL | 17 | 12,6 |
| CLL | 2 | 1,5 |
| HL | 4 | 3,0 |
| MPD | 15 | 11,1 |
| MM | 11 | 8,1 |
| Other | 6 | 4,4 |
| **Donor type** | | |
| Sibling | 79 | 58,5 |
| MUD | 56 | 41,5 |
| **Stem Cell Source** | | |
| Peripheral blood | 122 | 90,4 |
| Bone marrow | 13 | 9,6 |
| **HLA match**** | | |
| Match | 114 | 84,4 |
| Mismatch | 13 | 9,6 |
| Haplo | 7 | 5,2 |
| **Conditioning*** | | |
| MAC | 64 | 47,4 |
| RIC | 70 | 51,9 |
| **Prophylaxis of acute GVHD** | | |
| Calcineurin-based | 125 | 92,6 |
| mTOR-inhibitor based | 10 | 7,4 |
| **Acute GVHD**** | | |
| No | 66 | 48,9 |
| Grade <2 | 17 | 12,6 |
| Grade >2 | 50 | 37,0 |
| **Chronic GVHD*** | | |
| Mild | 28 | 20,7 |
| Moderate-Severe | 106 | 78,5 |
| **Delay from tx to sampling** | | |
| < 12 m | 67 | 49,6 |

| | | |
|---|---|---|
| 12-36 m | 53 | 39,3 |
| > 36 m | 15 | 11,1 |

**B. Patients without cGVHD**

| **Patient characteristics** | **N** | **%** |
|---|---|---|
| **Total** | 38 | |
| **Age at sampling (mean, range)** | 48 | (16-70) |
| **Patient sex** | | |
| Female | 23 | 60,5 |
| Male | 15 | 39,5 |
| **Donor sex*** | | |
| Female | 12 | 31,6 |
| Male | 26 | 68,4 |
| **Sex mismatch*** | | |
| No | 20 | 52,6 |
| Yes | 18 | 47,4 |
| **Diagnosis** | | |
| AML, MDS | 23 | 60,5 |
| ALL | 6 | 15,8 |
| NHL | 2 | 5,3 |
| CLL | 2 | 5,3 |
| HL | 3 | 7,9 |
| MPD | 1 | 2,6 |
| MM | 0 | 0 |
| Other | 1 | 2,6 |
| **Donor type** | | |
| Sibling | 23 | 60,5 |
| MUD | 15 | 39,5 |
| **Stem Cell Source** | | |
| Peripheral blood | 33 | 86,8 |
| Bone marrow | 5 | 13,2 |
| **HLA match*** | | |
| Match | 29 | 76,3 |
| Mismatch | 2 | 5,3 |
| Haplo | 6 | 15,8 |
| **Conditioning** | | |
| MAC | 20 | 52,6 |
| RIC | 18 | 47,4 |
| **Prophylaxis of acute GVHD** | | |
| Calcineurin-based | 37 | 97,4 |
| mTOR-inhibitor based | 1 | 2,6 |
| **Acute GVHD** | | |
| No | 18 | 47,4 |
| Grade <2 | 8 | 21,1 |
| Grade >2 | 12 | 31,5 |
| **Delay from tx to sampling** | | |
| < 12 m | 14 | 36,8 |
| 12-36 m | 22 | 57,9 |
| > 36 m | 2 | 5,3 |

AML; acute myeloid leukemia, MDS; myelodysplastic syndrome, NHL; non-hodgkin lymphoma, CLL; chronic lymphocytic leukemia, HL; Hodgkin's lymphoma, MPD; myeloproliferative syndrome, MM; multiple myeloma. MUD; match unrelated donor. Haplo; haploidentical HLA. MAC; myeloablative, RIC; reduced-intensity conditioning, tx; transplantation. *one missing **two missing

**Supplementary Table S2. Gene list in the immunogene panel sequencing. Gene names are presented as HUGO Gene Nomenclature Committee (HGNC) symbols.**

| | | | | | | | | | |
|---|---|---|---|---|---|---|---|---|---|
| A2M | A2ML1 | ABCB1 | ABCF1 | ACE | ADA | ADAM10 | ADAM17 | ADAM8 | AGFG1 |
| AICDA | AIMP1 | AIRE | ALCAM | ALK | ANP32B | ANPEP | ANXA6 | ATM | B2M |
| BACH2 | BANK1 | BATF | BAX | BCAM | BCAP31 | BCL2 | BCL2L1 | BCL6 | BIRC3 |
| BLK | BLM | BMP2 | BSG | BST1 | BST2 | BTK | BTLA | C1QA | C1QB |
| C1QBP | C1QC | C1QL1 | C1QL2 | C1QL3 | C1QL4 | C1QTNF2 | C1QTNF3 | C1QTNF4 | C1QTNF5 |
| C1QTNF6 | C1QTNF7 | C1R | C1RL | C1S | C2 | C3 | C3AR1 | C4A | C4B |
| C4BPA | C4BPB | C5 | C5AR1 | C6 | C7 | C8A | C8B | C8G | C9 |
| C9orf47 | CAMP | CANX | CASP1 | CASP10 | CASP2 | CASP3 | CASP4 | CASP6 | CASP7 |
| CASP8 | CASP9 | CCBP2 | CCL1 | CCL11 | CCL13 | CCL14 | CCL15 | CCL16 | CCL17 |
| CCL18 | CCL19 | CCL2 | CCL20 | CCL21 | CCL22 | CCL23 | CCL24 | CCL25 | CCL26 |
| CCL27 | CCL28 | CCL3 | CCL3L1 | CCL3L3 | CCL4 | CCL4L1 | CCL4L2 | CCL5 | CCL7 |
| CCL8 | CCND2 | CCND3 | CCNE1 | CCR1 | CCR10 | CCR2 | CCR3 | CCR4 | CCR5 |
| CCR6 | CCR6 | CCR7 | CCR9 | CCRL1 | CCRL2 | CCRN4L | CD101 | CD109 | CD14 |
| CD151 | CD160 | CD163 | CD164 | CD164L2 | CD180 | CD19 | CD1A | CD1B | CD1C |
| CD1D | CD1E | CD2 | CD200R1 | CD200R1L | CD207 | CD209 | CD22 | CD226 | CD244 |
| CD247 | CD248 | CD27 | CD274 | CD276 | CD28 | CD2AP | CD2BP2 | CD300A | CD300C |
| CD300E | CD300LB | CD300LF | CD300LG | CD302 | CD320 | CD33 | CD34 | CD36 | CD37 |
| CD38 | CD3D | CD3E | CD3EAP | CD3G | CD4 | CD40 | CD40LG | CD44 | CD46 |
| CD47 | CD48 | CD5 | CD52 | CD53 | CD58 | CD59 | CD5L | CD6 | CD63 |
| CD68 | CD69 | CD7 | CD70 | CD72 | CD74 | CD79A | CD79B | CD80 | CD81 |
| CD82 | CD83 | CD84 | CD86 | CD8A | CD8B | CD9 | CD93 | CD96 | CD97 |
| CD99 | CD99L2 | CDH5 | CDK6 | CDKN1A | CEACAM1 | CEACAM3 | CEACAM5 | CEACAM6 | CEACAM8 |
| CEBPB | CEBPE | CFD | CFH | CFHR1 | CFHR2 | CFHR3 | CFHR4 | CFHR5 | CFI |
| CFLAR | CFP | CHEK1 | CHL1 | CHUK | CIITA | CISH | CKLF | CLCF1 | CLEC10A |
| CLEC12A | CLEC16A | CLEC4A | CLEC4C | CLEC4D | CLEC4E | CLEC4M | CLEC5A | CLEC6A | CLEC7A |
| CLECL1 | CLIP1 | CLIP2 | CLU | CMKLR1 | CMTM1 | CMTM2 | CMTM3 | CMTM4 | CMTM5 |
| CMTM6 | CMTM7 | CMTM8 | COLEC12 | CR1 | CR1L | CR2 | CRADD | CRLF1 | CRLF2 |
| CRLF3 | CRP | CSF1 | CSF1R | CSF2 | CSF2RA | CSF2RB | CSF3 | CSF3R | CSN2 |
| CTLA4 | CTSG | CTSS | CX3CL1 | CX3CR1 | CXCL1 | CXCL10 | CXCL11 | CXCL12 | CXCL13 |
| CXCL14 | CXCL16 | CXCL2 | CXCL3 | CXCL5 | CXCL6 | CXCL9 | CXCR1 | CXCR2 | CXCR3 |
| CXCR4 | CXCR5 | CXCR6 | CXCR7 | CYBA | CYBB | CYSLTR1 | CYTL1 | DARC | DCD |
| DCLRE1C | DDR1 | DEFA1 | DEFA3 | DEFA4 | DEFA5 | DEFA6 | DEFB1 | DEFB103A | DEFB105A |
| DEFB106A | DEFB119 | DEFB123 | DEFB4A | DKC1 | DOCK2 | DPP4 | DUSP1 | EBF1 | EBF2 |
| EBI3 | EGF | EIF2AK2 | ELK1 | EMR3 | ENC1 | ENG | ENTPD1 | EPO | EPX |
| ERAP1 | ERGIC2 | ERLIN1 | ETS1 | ETS2 | F3 | FADD | FAK | FAS | FASLG |
| FCAMR | FCAR | FCER1A | FCER1G | FCER2 | FCGR1A | FCGR2A | FCGR2B | FCGR2C | FCGR3A |
| FCGR3B | FCGRT | FCRL5 | FCRLA | FGFR1 | FGFR2 | FGFR3 | FGFR4 | FGR | FLT3 |
| FOS | FOXK2 | FOXN1 | FOXO1 | FOXO3 | FOXP3 | FRK | FUT3 | FYN | G6PD |
| GADD45A | GNLY | GP1BA | GP1BB | GP5 | GP9 | GPR183 | GUSB | GYPA | GYPB |
| GYPC | GYPE | GZMA | GZMB | GZMK | GZMM | HAMP | HCK | HGF | HLA-A |
| HLA-B | HLA-C | HLA-DMA | HLA-DMB | HLA-DOA | HLA-DOB | HLA-DPA1 | HLA-DPB1 | HLA-DQA1 | HLA-DQB1 |
| HLA-DQB2 | HLA-DRA | HLA-DRB5 | HLA-E | HLA-F | HMGB1 | HMMR | HPS3 | HRAS | HRH2 |
| HRH4 | HSP90B1 | HSPA4 | HSPA6 | HSPD1 | HTN3 | ICAM1 | ICAM2 | ICAM3 | ICAM4 |
| ICOS | IFI16 | IFI27 | IFI35 | IFI44L | IFIH1 | IFIT1 | IFIT1B | IFIT2 | IFIT3 |
| IFITM1 | IFNA2 | IFNAR2 | IFNB1 | IFNG | IFNGR1 | IGF1R | IGF2R | IGJ | IGLL1 |
| IGSF8 | IKBKAP | IKBKB | IKBKE | IKBKG | IKZF1 | IKZF2 | IKZF3 | IL-17 | IL10 |
| IL10RA | IL10RB | IL11 | IL11RA | IL12A | IL12B | IL12B | IL12RB1 | IL12RB2 | IL13 |
| IL13RA1 | IL13RA2 | IL15 | IL15RA | IL16 | IL17A | IL17B | IL17C | IL17D | IL17F |
| IL17RA | IL17RB | IL17RC | IL17RD | IL17RE | IL18 | IL18BP | IL18R1 | IL18RAP | IL19 |
| IL1A | IL1B | IL1F10 | IL1R1 | IL1R2 | IL1RAP | IL1RAPL1 | IL1RAPL2 | IL1RL1 | IL1RL2 |
| IL1RN | IL2 | IL20 | IL20RA | IL21 | IL21R | IL22 | IL22RA1 | IL22RA2 | IL23A |
| IL23R | IL24 | IL25 | IL26 | IL27 | IL27RA | IL28A | IL28B | IL28RA | IL29 |
| IL2RA | IL2RB | IL2RG | IL3 | IL31 | IL31RA | IL32 | IL33 | IL36A | IL36G |

| | | | | | | | | | |
|---|---|---|---|---|---|---|---|---|---|
| IL36RN | IL37 | IL3RA | IL4 | IL4I1 | IL4R | IL5 | IL5RA | IL6 | IL6R |
| IL6ST | IL7 | IL7R | IL8 | IL9 | IL9R | ILF2 | ILF3 | INDO | INSR |
| IRAK1 | IRAK1BP1 | IRAK2 | IRAK3 | IRAK4 | IRF1 | IRF2 | IRF4 | IRF5 | IRF8 |
| IRF9 | ISG20 | ITFG1 | ITGA1 | ITGA2 | ITGA2B | ITGA3 | ITGA4 | ITGA5 | ITGA6 |
| ITGAD | ITGAE | ITGAL | ITGAM | ITGAV | ITGAX | ITGB1 | ITGB2 | ITGB3 | ITGB4 |
| JAK1 | JAK2 | JAK3 | JUN | KDR | KEL | KIF21B | KIR2DL1 | KIR2DL3 | KIR2DL4 |
| KIR2DS4 | KIR3DL1 | KIR3DL2 | KIR3DL3 | KIT | KITLG | KLRB1 | KLRC1 | KLRC2 | KLRD1 |
| KLRK1 | KRAS | L1CAM | LAG3 | LAIR1 | LAIR2 | LAMP1 | LAMP2 | LAMP3 | LAMTOR3 |
| LAX1 | LCK | LCP2 | LEAP2 | LEP | LIF | LIFR | LIG1 | LIG4 | LILRA1 |
| LILRA2 | LILRA3 | LILRA4 | LILRA4 | LILRA5 | LILRA6 | LILRB1 | LILRB2 | LILRB3 | LILRB4 |
| LILRB5 | LITAF | LPO | LRP1 | LTA | LTB | LTB4R | LTB4R2 | LTBR | LTF |
| LY75 | LY86 | LY9 | LY96 | LYG2 | LYN | LYZ | MAF | MAL | MAP2K3 |
| MAP2K4 | MAP2K6 | MAP3K1 | MAP3K14 | MAP4K4 | MAPK1 | MAPK10 | MAPK11 | MAPK12 | MAPK13 |
| MAPK14 | MAPK3 | MAPK6 | MAPK7 | MAPK8 | MAPK8IP3 | MAPK9 | MAPKAPK2 | MARCO | MASP1 |
| MASP2 | MBL2 | MBP | MBTPS1 | MCAM | MCL1 | MDM2 | MFI2 | MICA | MICB |
| MIF | MME | MMP9 | MPL | MPO | MR1 | MRC1 | MRC2 | MRE11A | MS4A1 |
| MS4A3 | MS4A5 | MSR1 | MST1 | MST1R | MTOR | MUC1 | MX1 | MYC | MYD88 |
| MYLK | NCAM1 | NCF1 | NCF2 | NCF4 | NCR1 | NCR2 | NCR3 | NDUFS3 | NFATC1 |
| NFATC2 | NFATC3 | NFATC4 | NFIL3 | NFKB1 | NFKB2 | NFKBIA | NFKBIB | NFKBIE | NFKBIL1 |
| NFRKB | NKX2-3 | NOD2 | NOL3 | NOP9 | NOS2 | NPTN | NRAS | NT5E | PADI4 |
| PAFAH1B1 | PAFAH1B2 | PAFAH1B3 | PAFAH2 | PARP1 | PDCD1 | PDCD1LG2 | PDGFB | PDGFRA | PDGFRB |
| PELI1 | PELI2 | PF4 | PGLYRP1 | PGLYRP2 | PGLYRP3 | PGLYRP4 | PIAS3 | PIK3CG | PIK3R1 |
| PILRA | PIM1 | PLA2G7 | PLA2R1 | PLAA | PLAUR | PLK3 | PLXNC1 | PNP | POMC |
| POU2AF1 | PPBP | PPIA | PPP3CA | PPP3CB | PPP3CC | PPP3R1 | PPP3R2 | PRDM1 | PRDX6 |
| PRF1 | PRG2 | PRKCD | PRKCQ | PRNP | PROCR | PROM1 | PRSS16 | PSG1 | PSIP1 |
| PSMB10 | PSMB5 | PSMB6 | PSMB7 | PSMB8 | PSMB9 | PSME1 | PSME2 | PSME3 | PSMF1 |
| PSTPIP1 | PTAFR | PTEN | PTGER4 | PTGFRN | PTGS2 | PTK2 | PTK2B | PTPN11 | PTPN2 |
| PTPN22 | PTPN6 | PTPRC | PTPRCAP | PTPRE | PTPRJ | PVR | PVRL1 | PVRL2 | PXK |
| RAC1 | RAC2 | RAG1 | RAG2 | RBPJ | REL | RELA | RELB | RFX1 | RFX5 |
| RFXANK | RFXAP | RGS1 | RHAG | RHCE | RHD | RIPK1 | RIPK2 | RNASE7 | ROR1 |
| RORA | RORC | RPA1 | S100A8 | SAMSN1 | SARM1 | SCARB1 | SCARB2 | SCGB3A1 | SDC1 |
| SDF2 | SDF2L1 | SELE | SELL | SELP | SELPLG | SEMA4D | SEMA7A | SERPING1 | SH2B3 |
| SH2D1A | SIGIRR | SIGLEC1 | SIGLEC15 | SIGLEC5 | SIGLEC6 | SIRPA | SIVA1 | SLA | SLAMF1 |
| SLAMF6 | SLAMF7 | SLC3A2 | SLC44A1 | SLC4A1 | SLC7A5 | SMARCAL1 | SOCS1 | SOCS2 | SOCS3 |
| SOCS4 | SOCS5 | SOCS6 | SOCS7 | SOD1 | SOD3 | SPN | SPP1 | SRC | ST6GAL1 |
| STAT1 | STAT2 | STAT3 | STAT4 | STAT5A | STAT5B | STAT6 | TAGAP | TAL1 | TANK |
| TAP1 | TAP2 | TAP2 | TAPBP | TBK1 | TBX21 | TCF3 | TCF7 | TCN2 | TDP2 |
| TEK | TFRC | TGFB1 | THBD | THY1 | TICAM1 | TICAM2 | TIMP1 | TIRAP | TLR1 |
| TLR10 | TLR2 | TLR3 | TLR4 | TLR5 | TLR6 | TLR7 | TLR8 | TLR9 | TLR9 |
| TMED7 | TNF | TNFAIP3 | TNFRSF10A | TNFRSF10B | TNFRSF10C | TNFRSF10D | TNFRSF11A | TNFRSF12A | TNFRSF13B |
| TNFRSF13B | TNFRSF13C | TNFRSF14 | TNFRSF17 | TNFRSF18 | TNFRSF1A | TNFRSF1B | TNFRSF25 | TNFRSF4 | TNFRSF8 |
| TNFRSF9 | TNFSF10 | TNFSF11 | TNFSF12 | TNFSF13 | TNFSF13B | TNFSF14 | TNFSF15 | TNFSF4 | TNFSF8 |
| TNIP1 | TOLLIP | TONSL | TRADD | TRAF1 | TRAF2 | TRAF3 | TRAF3IP1 | TRAF4 | TRAF5 |
| TRAF6 | TRAF7 | TSPYL2 | TYK2 | ULBP1 | UNG | WAS | WASF1 | WASF3 | VCAM1 |
| VCAN | VEGFA | WFDC12 | WIPF1 | VPREB1 | XCL1 | XCL2 | XCR1 | XRCC5 | YES |
| YWHAZ | ZAP70 | ZEB1 | ZFP36 | ZFP36L1 | | | | | |

**Supplementary Table S3. Primer sets of mTOR, NFkB2 and TLR2 amplicon sequencing**

| Primer name | Sequence |
|---|---|
| mTOR-F | 5’-ACACTCTTTCCCTACACGACGCTCTTCCGATCTTCCCTGTAGTCCCGGATGAG-3’ |
| mTOR-R | 5’-AGACGTGTGCTCTTCCGATCTGCCTGTGTTCTGAGCTGCTC-3’ |
| NFkB2-F | 5’-ACACTCTTTCCCTACACGACGCTCTTCCGATCTATCCCATTCCTGTCCCCATTT-3’ |
| NFkB2-R | 5’-AGACGTGTGCTCTTCCGATCTAGTGCACCTGAGGCTGGG-3’ |
| TLR2-F | 5’-ACACTCTTTCCCTACACGACGCTCTTCCGATCTAAGACTTTGGAAGCTGGTGG-3’ |
| TLR2-R | 5’-AGACGTGTGCTCTTCCGATCTCGGACATCCTGAACCTGCT-3’ |

Abbreviations: F, forward primer; R, reverse primer

**Supplementary Table S4. Primer set for mTOR, NFkB2 and TLR2 capillary sequencing**

| Gene | Forward primer | Reverse primer | Product length |
|---|---|---|---|
| MTOR | 5’-TCTGCCTGTGTTCTGAGCTG-3’ | 5’-CGATGCTCGATGTTGAGAAG-3’ | 210 |
| NFkB2 | 5’-CAACTCCGGATCTCGCTCTC-3’ | 5’-GGCATGACTCACTGGGTTGT-3’ | 259 |
| TLR2 | 5’-GCCTCCCTCTTACCCATGTT-3’ | 5’-TACCACAGGCCATGGAAACG-3’ | 368 |

**Supplementary Table S5. List of mTOR, NFkB2 and TLR2 mutagenesis primers**

| Primer name | Sequence |
|---|---|
| mTOR_P2229R-F | 5’-GAGATACGCTGTCATCCGTTTATCGACCAACTCGG-3’ |
| mTOR_P2229R-R | 5’-CCGAGTTGGTCGATAAACGGATGACAGCGTATCTC-3’ |
| NFkB2_P882Q-F | 5’-TCCCCCAAATCTGGGCCACCATCTCCCGCCCACCC-3’ |
| NFkB2_P882Q-R | 5’-GGGTGGGCGGGAGATGGTGGCCCAGATTTGGGGGA-3’ |
| TLR2_W558L-F | 5’-CAAAGTCTTGATTGATTTGCCAGCAAATTACCTGT-3’ |
| TLR2_W558L-R | 5’-ACAGGTAATTTGCTGGCAAATCAATCAAGACTTTG-3’ |

Abbreviations: F, forward primer; R, reverse primer

**Supplementary Table S6. Primer list of RT-qPCR**

| Target gene | Sequence |
|---|---|
| FOS-F | 5’-CCGGGGATAGCCTCTCTTACT-3’ |
| FOS-R | 5’-CCAGGTCCGTGCAGAAGTC-3’ |
| ELK1-F | 5’-AATCGGAAGAGCTTAATGTGGAG-3’ |
| ELK1-R | 5’-CTTGGTGGTTTCTGGCACAA-3’ |
| ACTB-F | 5’-GTTGTCGACGACGAGCG-3’ |
| ACTB-R | 5’-GCACAGAGCCTCGCCTT-3’ |

Abbreviations: F, forward primer; R, reverse primer

**Supplementary Table S7. Immunogene panel sequencing result for the index patient**

(A) CD8+ T-cells purified in 2013

| Chr | position | ref | var | Gene | Mutation type | Codon change | Exon | Amino acid change | Normal_ reads [1] | Normal_ reads [2] | Normal_var_ Freq (%) | Tumor_ reads [3] | Tumor_ reads [4] | Tumor_var_ Freq (%) | Somatic p-valu* |
|---|---|---|---|---|---|---|---|---|---|---|---|---|---|---|---|
| chr17 | 80274141 | GCAGGGA GGGCAGA GGCTGCT GGCGGGT | G | CD7 | CODON CHANGE PLUS DELETION | gacccgccag cagcctctgcc ctccctgcg/ gcg | 3 | DPPAASALPA1 72A | 19 | 0 | 0 | 8 | 7 | 46.67 | 0.0011962 |

(B) CD4+ T-cells purified in 2015

| Chr | position | ref | var | Gene | Mutation type | Codon change | Exon | Amino acid change | Normal_ reads [1] | Normal_ reads [2] | Normal_var_ Freq (%) | Tumor_ reads [3] | Tumor_ reads [4] | Tumor_var_ Freq (%) | Somatic p-valu* |
|---|---|---|---|---|---|---|---|---|---|---|---|---|---|---|---|
| 22 | 29885598 | AAGGAAG | A | NEFH | CODON_DELETION | aaggaagag/ aag | 4 | KEE657K | 141 | 0 | 0 | 114 | 21 | 15.56 | 1.2713E-07 |
| 22 | 30198134 | A | T | ASCC2 | MISSENSE | Tcc/Acc | 14 | S473T | 48 | 0 | 0 | 28 | 13 | 31.71 | 0.000012532 |
| 11 | 45671492 | C | T | CHST1 | MISSENSE | Gtg/Atg | 4 | V328M | 76 | 0 | 0 | 48 | 11 | 18.64 | 0.000062539 |
| 17 | 8160230 | C | A | PFAS | MISSENSE | Cac/Aac | 9 | H342N | 48 | 0 | 0 | 26 | 10 | 27.78 | 0.000092062 |
| 1 | 236906279 | G | T | ACTN2 | MISSENSE | gaG/gaT | 11 | E397D | 91 | 0 | 0 | 72 | 11 | 13.25 | 0.000201 |
| 4 | 100573255 | C | T | RP11-766 F14.2.1 | MISSENSE | Gcc/Acc | 1 | A851T | 83 | 0 | 0 | 56 | 9 | 13.85 | 0.00043634 |
| 17 | 66986988 | T | C | ABCA9 | MISSENSE | gAc/gGc | 29 | D1276G | 44 | 0 | 0 | 18 | 7 | 28 | 0.00044555 |
| 12 | 6181540 | G | A | VWF | MISSENSE | Cgc/Tgc | 9 | R356C | 56 | 0 | 0 | 45 | 10 | 18.18 | 0.00056748 |
| **10** | **104162075** | **C** | **A** | **NFKB2** | **MISSENSE** | **cCa/cAa** | **23** | **P882Q** | **113** | **0** | **0** | **75** | **8** | **9.64** | **0.0008439** |
| 3 | 119886213 | C | T | GPR156 | MISSENSE | cGg/cAg | 10 | R704Q | 58 | 0 | 0 | 37 | 8 | 17.78 | 0.00090659 |
| 22 | 30927925 | C | T | SEC14L6 | MISSENSE | Gat/Aat | 6 | D162N | 66 | 0 | 0 | 43 | 8 | 15.69 | 0.00093377 |
| 9 | 34834390 | C | A | AL589645. 1 | MISSENSE | Gtg/Ttg | 4 | V367L | 138 | 0 | 0 | 113 | 9 | 7.38 | 0.00093765 |
| **1** | **11182160** | **G** | **C** | **MTOR** | **MISSENSE** | **cCt/cGt** | **48** | **P2229R** | **55** | **0** | **0** | **23** | **6** | **20.69** | **0.0011686** |
| **4** | **154625732** | **G** | **T** | **TLR2** | **MISSENSE** | **tGg/tTg** | **1** | **W558L** | **63** | **0** | **0** | **54** | **7** | **11.48** | **0.0057957** |
| 2 | 44428602 | TGGAAAA TCAG | T | PPM1B | FRAME_SHIFT | -/- | 2 | -89 | 88 | 0 | 0 | 63 | 6 | 8.7 | 0.0063483 |
| 17 | 72926604 | G | T | OTOP2 | MISSENSE | Ggc/Tgc | 6 | G292C | 85 | 0 | 0 | 65 | 6 | 8.45 | 0.0078858 |
| 7 | 141401764 | G | A | KIAA1147 | MISSENSE | Ccc/Tcc | 1 | P64S | 16 | 0 | 0 | 5 | 4 | 44.44 | 0.0099605 |

### (C) CD8+ T-cells purified in 2015

| Chr | position | ref | var | Gene | Mutation type | Codon change | Ex on | Amino acid change | Normal_ reads [1] | Normal_ reads [2] | Normal_var_ Freq (%) | Tumor_ reads [3] | Tumor_ reads [4] | Tumor_var_ Freq (%) | Somatic p-valu* |
|---|---|---|---|---|---|---|---|---|---|---|---|---|---|---|---|
| chr22 | 26879946 | GGAGGCG GCGCCCC GGGGGA GA | G | SRRD | DELETION | gaggcggcgcccc gggggaga/- | 1 | EAAPRGR31- | 10 | 0 | 0 | 6 | 7 | 53.85 | 0.0069996 |
| chr21 | 47612645 | G | C | AP001468.1 | MISSENSE | cCc/cGc | 2 | P223R | 32 | 1 | 3.03 | 30 | 10 | 25 | 0.0085105 |
| chr12 | 53069222 | CACCTCC GGAGCCG TAGCTGC T | C | KRT1 | DELETION | agcagctacggct ccggaggt/- | 9 | SSYGSGG557- | 8 | 0 | 0 | 13 | 14 | 51.85 | 0.008646 |

### (D) NK cells purified in 2015

| Chr | position | ref | var | Gene | Mutation type | Codon change | Ex on | Amino acid change | Normal_ reads [1] | Normal_ reads [2] | Normal_var_ Freq (%) | Tumor_ reads [3] | Tumor_ reads [4] | Tumor_var_ Freq (%) | Somatic p-valu* |
|---|---|---|---|---|---|---|---|---|---|---|---|---|---|---|---|
| chr12 | 21997770 | C | T | ABCC9 | STOP_GAINED | tGg/tAg | 25 | W1059* | 70 | 0 | 0 | 78 | 32 | 29.09 | 1.7846E-08 |
| chr18 | 28935163 | G | A | DSG1 | MISSENSE | Ggc/Agc | 15 | G1002S | 60 | 0 | 0 | 69 | 14 | 16.87 | 0.00029757 |
| chr1 | 231830034 | T | A | DISC1 | MISSENSE | cTg/cAg | 2 | L177Q | 81 | 0 | 0 | 72 | 10 | 12.2 | 0.00077705 |
| chr1 | 75606686 | G | A | LHX8 | STOP_GAINED | tGg/tAg | 5 | W95* | 37 | 0 | 0 | 34 | 10 | 22.73 | 0.0013209 |
| chr2 | 233839411 | T | C | NGEF | MISSENSE | Atc/Gtc | 2 | I64V | 52 | 0 | 0 | 51 | 10 | 16.39 | 0.0014524 |
| chr1 | 89449434 | T | C | RBMXL1 | MISSENSE | Aca/Gca | 2 | T26A | 28 | 1 | 3.45 | 19 | 10 | 34.48 | 0.0027031 |
| chr6 | 43266374 | C | T | SLC22A7 | MISSENSE | aCg/aTg | 1 | T93M | 28 | 0 | 0 | 16 | 6 | 27.27 | 0.0046954 |

Abbreviations: Chr, chromosome; ref, reference base; var, variant base; freq, frequency
1 Sequencing reads supporting reference allele in normal sample
2 Sequencing reads supporting variant allele in normal sample
3 Sequencing reads supporting reference allele in tumor sample
4 Sequencing reads supporting variant allele in tumor sample
* Somatic p-value for somatic/loss of heterozygosity events

**Supplementary Table S8. Somatic NFKB2, TLR2 and MTOR mutations in different cellular fractions (Supplementary Figure S2C) from the index patient validated by amplicon sequencing**

| DNAs | Gene | Chr | position | ref | var | Amino acid change | Total_Depth | Call_Depth | Ref_Calls | Var_Calls | VAF (%) | Freq_Ratio |
|---|---|---|---|---|---|---|---|---|---|---|---|---|
| CD3neg | NFKB2 | 10 | 104162075 | C | A | P882Q | 49669 | 49669 | 49640 | 29 | **0** | 0 |
| CD3neg | TLR2 | 4 | 154625732 | G | T | W558L | 102539 | 102539 | 102411 | 128 | **0** | 0 |
| CD3neg | MTOR | 1 | 11182160 | G | C | P2229R | 133746 | 133746 | 133628 | 118 | **0** | 0 |
| CD8+ | NFKB2 | 10 | 104162075 | C | A | P882Q | 13945 | 13944 | 13936 | 8 | **0** | 0 |
| CD8+ | TLR2 | 4 | 154625732 | G | T | W558L | 125028 | 125028 | 124912 | 116 | **0** | 0 |
| CD8+ | MTOR | 1 | 11182160 | G | C | P2229R | 156055 | 156055 | 155995 | 60 | **0** | 0 |
| CD4+Vb.20neg | NFKB2 | 10 | 104162075 | C | A | P882Q | 21047 | 21047 | 21024 | 23 | **0** | 0 |
| CD4+Vb.20neg | TLR2 | 4 | 154625732 | G | T | W558L | 149991 | 149991 | 149151 | 840 | **0.5** | 0.98095 |
| CD4+Vb.20neg | MTOR | 1 | 11182160 | G | C | P2229R | 184213 | 184213 | 183230 | 983 | **0** | 0 |
| CD4+CD8+ Vb.20neg | NFKB2 | 10 | 104162075 | C | A | P882Q | 66813 | 66812 | 66301 | 511 | **0.7** | 0.98661 |
| CD4+CD8+ Vb.20neg | TLR2 | 4 | 154625732 | G | T | W558L | 124985 | 124985 | 123059 | 1926 | **1.5** | 1.00066 |
| CD4+CD8+ Vb.20neg | MTOR | 1 | 11182160 | G | C | P2229R | 170803 | 170803 | 167577 | 3226 | **1.9** | 0.99731 |
| Monocytes | NFKB2 | 10 | 104162075 | C | A | P882Q | 40654 | 40653 | 40618 | 35 | **0** | 0 |
| Monocytes | TLR2 | 4 | 154625732 | G | T | W558L | 147684 | 147684 | 146326 | 1358 | **0.9** | 0.99539 |
| Monocytes | MTOR | 1 | 11182160 | G | C | P2229R | 172128 | 172128 | 169802 | 2326 | **1.3** | 0.99164 |

The purity of sorted fractions was evaluated by flow cytometry (FACs AriaII) and confirmed to be >98%

Abbreviations: Chr, chromosome; ref, reference base; var, variant base; VAF, variant allele frequency; Freq, frequency; Freq_Ratio, base quality frequency ratio

**Supplementary Table S9. Somatic TLR2 mutation in CD4+ T-cells validated by amplicon sequencing**

| | Patient | DNAs | Gene | Chr | position | ref | var | Amino acid change | Total_Depth | Call_Depth | Ref_Calls | Var_Calls | VAF (%) | Freq_Ratio |
|---|---|---|---|---|---|---|---|---|---|---|---|---|---|---|
| 1-1 | GVHD | CD4+T | TLR2 | 4 | 154625732 | G | T | W558L | 543370 | 543370 | 453452 | 89918 | 16.529 | 1.00623 |
| 1-2 | GVHD | CD4+Vb.20 | TLR2 | 4 | 154625732 | G | T | W558L | 160135 | 160135 | 103031 | 57104 | 35.521 | 1.01047 |
| 1-3 | GVHD | CD4+CD8+ Vb.20 | TLR2 | 4 | 154625732 | G | T | W558L | 123431 | 123431 | 89029 | 34402 | 27.838 | 1.01117 |
| 2 | GVHD | CD4+T | TLR2 | 4 | 154625732 | G | T | W558L | 39775 | 39775 | 34242 | 5533 | 13.863 | 1.01118 |
| 3 | GVHD | CD4+T | TLR2 | 4 | 154625732 | G | T | W558L | 31285 | 31285 | 26791 | 4494 | 14.31 | 1.01125 |
| 4 | GVHD | CD4+T | TLR2 | 4 | 154625732 | G | T | W558L | 17162 | 17162 | 15779 | 1383 | 8.029 | 1.01096 |
| 5 | GVHD | CD4+T | TLR2 | 4 | 154625732 | G | T | W558L | 31044 | 31044 | 27152 | 3892 | 12.492 | 1.01145 |
| 6 | GVHD | CD4+T | TLR2 | 4 | 154625732 | G | T | W558L | 15876 | 15876 | 15067 | 809 | 5.058 | 1.00573 |
| 7 | GVHD | CD4+T | TLR2 | 4 | 154625732 | G | T | W558L | 28823 | 28823 | 26310 | 2513 | 8.67 | 1.00554 |
| 8 | GVHD | CD4+T | TLR2 | 4 | 154625732 | G | T | W558L | 16932 | 16932 | 15291 | 1641 | 9.68 | 1.0061 |
| 9 | GVHD | CD4+T | TLR2 | 4 | 154625732 | G | T | W558L | 19309 | 19309 | 17983 | 1326 | 6.836 | 1.0079 |
| 10 | GVHD | CD4+T | TLR2 | 4 | 154625732 | G | T | W558L | 28042 | 28042 | 25247 | 2795 | 9.949 | 1.00633 |
| 11 | GVHD | CD4+T | TLR2 | 4 | 154625732 | G | T | W558L | 33628 | 33628 | 28176 | 5452 | 16.168 | 1.00513 |
| 12 | GVHD | CD4+T | TLR2 | 4 | 154625732 | G | T | W558L | 34034 | 34034 | 31716 | 2318 | 6.743 | 1.00415 |
| 13 | GVHD | CD4+T | TLR2 | 4 | 154625732 | G | T | W558L | 27185 | 27185 | 24024 | 3161 | 11.602 | 1.01008 |
| 14 | GVHD | CD4+T | TLR2 | 4 | 154625732 | G | T | W558L | 8934 | 8934 | 8290 | 644 | 7.152 | 1.00475 |
| 15 | GVHD | CD4+T | TLR2 | 4 | 154625732 | G | T | W558L | 25657 | 25657 | 21757 | 3900 | 15.165 | 1.01141 |
| 16 | GVHD | CD4+T | TLR2 | 4 | 154625732 | G | T | W558L | 24821 | 24821 | 23284 | 1537 | 6.156 | 1.00601 |
| 17 | GVHD | CD4+T | TLR2 | 4 | 154625732 | G | T | W558L | 32618 | 32617 | 29085 | 3532 | 10.764 | 1.01143 |
| 18 | GVHD | CD4+T | TLR2 | 4 | 154625732 | G | T | W558L | 20425 | 20425 | 19223 | 1202 | 5.821 | 1.01546 |
| 19 | GVHD | CD4+T | TLR2 | 4 | 154625732 | G | T | W558L | 24791 | 24791 | 20940 | 3851 | 15.485 | 1.00852 |
| 20 | GVHD | CD4+T | TLR2 | 4 | 154625732 | G | T | W558L | 32973 | 32973 | 30111 | 2862 | 8.646 | 1.00359 |
| 21 | GVHD | CD4+T | TLR2 | 4 | 154625732 | G | T | W558L | 456 | 456 | 379 | 77 | 16.667 | 1.02202 |
| 22 | control | CD4+T | TLR2 | 4 | 154625732 | G | T | W558L | 32297 | 32297 | 32066 | 231 | 0.666 | 0.97297 |
| 23 | control | CD4+T | TLR2 | 4 | 154625732 | G | T | W558L | 17599 | 17599 | 17490 | 109 | 0.563 | 0.97691 |
| 24 | control | CD4+T | TLR2 | 4 | 154625732 | G | T | W558L | 28586 | 28586 | 28296 | 290 | 0.983 | 0.98678 |
| 25 | control | CD4+T | TLR2 | 4 | 154625732 | G | T | W558L | 35632 | 35632 | 34855 | 777 | 2.133 | 1.00516 |
| 26 | control | CD4+T | TLR2 | 4 | 154625732 | G | T | W558L | 22900 | 22900 | 22428 | 472 | 1.983 | 0.99899 |

Abbreviations: Chr, chromosome; ref, reference base; var, variant base; VAF, variant allele frequency; Freq, frequency; Freq_Ratio, base quality frequency ratio

**Supplementary Table S10. Somatic TLR2 mutation in CD8+ T-cells validated by amplicon sequencing**

| | Patient | DNAs | Gene | Chr | position | ref | var | Amino acid change | Total_Depth | Call_Depth | Ref_Calls | Var_Calls | VAF (%) | Freq_Ratio |
|---|---|---|---|---|---|---|---|---|---|---|---|---|---|---|
| 1 | GVHD | CD8+T | TLR2 | 4 | 154625732 | G | T | W558L | 999903 | 999903 | 952631 | 47272 | 4.701 | 1.00787 |
| 2 | GVHD | CD8+T | TLR2 | 4 | 154625732 | G | T | W558L | 12699 | 12699 | 11570 | 1129 | 8.827 | 1.00725 |
| 3 | GVHD | CD8+T | TLR2 | 4 | 154625732 | G | T | W558L | 18488 | 18488 | 16762 | 1726 | 9.233 | 1.0104 |
| 4 | GVHD | CD8+T | TLR2 | 4 | 154625732 | G | T | W558L | 40286 | 40286 | 34158 | 6128 | 15.181 | 1.01074 |
| 5 | GVHD | CD8+T | TLR2 | 4 | 154625732 | G | T | W558L | 6416 | 6416 | 5960 | 456 | 7.014 | 1.01526 |
| 6 | GVHD | CD8+T | TLR2 | 4 | 154625732 | G | T | W558L | 54276 | 54276 | 51710 | 2566 | 4.656 | 1.00365 |
| 7 | GVHD | CD8+T | TLR2 | 4 | 154625732 | G | T | W558L | 36682 | 36682 | 33917 | 2765 | 7.51 | 1.01079 |
| 8 | GVHD | CD8+T | TLR2 | 4 | 154625732 | G | T | W558L | 15920 | 15920 | 14589 | 1331 | 8.285 | 1.00579 |
| 9 | GVHD | CD8+T | TLR2 | 4 | 154625732 | G | T | W558L | 19640 | 19640 | 17092 | 2548 | 12.943 | 1.01113 |
| 10 | GVHD | CD8+T | TLR2 | 4 | 154625732 | G | T | W558L | 37171 | 37171 | 34262 | 2909 | 7.791 | 1.00821 |
| 11 | GVHD | CD8+T | TLR2 | 4 | 154625732 | G | T | W558L | 24258 | 24258 | 21334 | 2924 | 12.008 | 1.00883 |
| 12 | GVHD | CD8+T | TLR2 | 4 | 154625732 | G | T | W558L | 29187 | 29187 | 26608 | 2579 | 8.795 | 1.0008 |
| 13 | GVHD | CD8+T | TLR2 | 4 | 154625732 | G | T | W558L | 26416 | 26416 | 25356 | 1060 | 3.967 | 0.99672 |
| 14 | GVHD | CD8+T | TLR2 | 4 | 154625732 | G | T | W558L | 20640 | 20640 | 19840 | 800 | 3.832 | 1.00522 |
| 15 | GVHD | CD8+T | TLR2 | 4 | 154625732 | G | T | W558L | 35883 | 35883 | 32070 | 3813 | 10.596 | 1.00623 |
| 16 | GVHD | CD8+T | TLR2 | 4 | 154625732 | G | T | W558L | 42754 | 42754 | 36774 | 5980 | 13.952 | 1.01269 |
| 17 | GVHD | CD8+T | TLR2 | 4 | 154625732 | G | T | W558L | 23141 | 23141 | 20973 | 2168 | 9.325 | 1.00794 |
| 18 | GVHD | CD8+T | TLR2 | 4 | 154625732 | G | T | W558L | 29169 | 29169 | 26838 | 2331 | 7.95 | 1.01069 |
| 19 | GVHD | CD8+T | TLR2 | 4 | 154625732 | G | T | W558L | 17956 | 17956 | 16985 | 971 | 5.324 | 1.01146 |
| 20 | GVHD | CD8+T | TLR2 | 4 | 154625732 | G | T | W558L | 35087 | 35087 | 31556 | 3531 | 10.032 | 1.00877 |
| 21 | GVHD | CD8+T | TLR2 | 4 | 154625732 | G | T | W558L | 20298 | 20298 | 19639 | 659 | 3.183 | 1.00346 |
| 22 | GVHD | CD8+T | TLR2 | 4 | 154625732 | G | T | W558L | 1704 | 1704 | 1598 | 106 | 6.045 | 0.99851 |
| 23 | control | CD8+T | TLR2 | 4 | 154625732 | G | T | W558L | 10509 | 10509 | 10354 | 155 | 1.418 | 1.00353 |
| 24 | control | CD8+T | TLR2 | 4 | 154625732 | G | T | W558L | 17666 | 17666 | 17327 | 339 | 1.834 | 0.99182 |
| 25 | control | CD8+T | TLR2 | 4 | 154625732 | G | T | W558L | 38778 | 38778 | 37897 | 881 | 2.192 | 1.01323 |
| 26 | control | CD8+T | TLR2 | 4 | 154625732 | G | T | W558L | 19211 | 19211 | 18069 | 1142 | 5.877 | 1.01361 |

Abbreviations: Chr, chromosome; ref, reference base; var, variant base; VAF, variant allele frequency; Freq, frequency; Freq_Ratio, base quality frequency ratio

### Supplementary Table S11. Gene Sets Enriched Analysis (GSEA)

| | GS follow link to MSigDB | GS DETAILS | SIZE | ES | NES | NOM p-val | FDR q-val | FWER p-val | RANK AT MAX | LEADING EDGE |
|---|---|---|---|---|---|---|---|---|---|---|
| 1 | HALLMARK_HYPOXIA | Details ... | 67 | 0.56 | 2.60 | 0.000 | 0.000 | 0.000 | 646 | tags=36%, list=12%, signal=40% |
| 2 | HALLMARK_MYC_TARGETS_V1 | Details ... | 128 | 0.51 | 2.57 | 0.000 | 0.000 | 0.000 | 730 | tags=40%, list=14%, signal=45% |
| 3 | HALLMARK_TNFA_SIGNALING_VIA_NFKB | Details ... | 87 | 0.52 | 2.41 | 0.000 | 0.000 | 0.000 | 734 | tags=44%, list=14%, signal=50% |
| 4 | HALLMARK_UV_RESPONSE_UP | Details ... | 63 | 0.47 | 2.14 | 0.000 | 0.002 | 0.004 | 519 | tags=27%, list=10%, signal=30% |
| 5 | HALLMARK_MTORC1_SIGNALING | Details ... | 100 | 0.41 | 1.96 | 0.000 | 0.013 | 0.030 | 625 | tags=29%, list=12%, signal=32% |
| 6 | HALLMARK_INTERFERON_GAMMA_RESPONSE | Details ... | 109 | 0.40 | 1.96 | 0.004 | 0.012 | 0.034 | 519 | tags=31%, list=10%, signal=34% |
| 7 | HALLMARK_MYC_TARGETS_V2 | Details ... | 32 | 0.50 | 1.88 | 0.008 | 0.018 | 0.064 | 730 | tags=28%, list=14%, signal=32% |
| 8 | HALLMARK_IL2_STAT5_SIGNALING | Details ... | 87 | 0.39 | 1.84 | 0.003 | 0.022 | 0.086 | 594 | tags=31%, list=11%, signal=34% |
| 9 | HALLMARK_GLYCOLYSIS | Details ... | 65 | 0.39 | 1.75 | 0.003 | 0.036 | 0.158 | 873 | tags=38%, list=17%, signal=46% |
| 10 | HALLMARK_ESTROGEN_RESPONSE_LATE | Details ... | 55 | 0.39 | 1.72 | 0.010 | 0.039 | 0.183 | 330 | tags=24%, list=6%, signal=25% |
| 11 | HALLMARK_ADIPOGENESIS | Details ... | 81 | 0.35 | 1.70 | 0.013 | 0.044 | 0.220 | 1004 | tags=38%, list=19%, signal=47% |
| 12 | HALLMARK_OXIDATIVE_PHOSPHORYLATION | Details ... | 118 | 0.31 | 1.59 | 0.007 | 0.082 | 0.391 | 796 | tags=32%, list=15%, signal=37% |
| 13 | HALLMARK_ESTROGEN_RESPONSE_EARLY | Details ... | 57 | 0.34 | 1.50 | 0.063 | 0.125 | 0.560 | 330 | tags=19%, list=6%, signal=20% |
| 14 | HALLMARK_PI3K_AKT_MTOR_SIGNALING | Details ... | 55 | 0.33 | 1.48 | 0.036 | 0.127 | 0.591 | 504 | tags=24%, list=10%, signal=26% |
| 15 | HALLMARK_IL6_JAK_STAT3_SIGNALING | Details ... | 31 | 0.39 | 1.47 | 0.083 | 0.129 | 0.623 | 243 | tags=16%, list=5%, signal=17% |
| 16 | HALLMARK_UNFOLDED_PROTEIN_RESPONSE | Details ... | 63 | 0.30 | 1.35 | 0.082 | 0.218 | 0.853 | 598 | tags=24%, list=11%, signal=27% |
| 17 | HALLMARK_REACTIVE_OXIGEN_SPECIES_PATHWAY | Details ... | 26 | 0.37 | 1.35 | 0.130 | 0.206 | 0.856 | 617 | tags=31%, list=12%, signal=35% |
| 18 | HALLMARK_KRAS_SIGNALING_UP | Details ... | 48 | 0.32 | 1.32 | 0.119 | 0.228 | 0.893 | 485 | tags=25%, list=9%, signal=27% |
| 19 | HALLMARK_HEME_METABOLISM | Details ... | 73 | 0.28 | 1.31 | 0.097 | 0.225 | 0.906 | 273 | tags=10%, list=5%, signal=10% |
| 20 | HALLMARK_FATTY_ACID_METABOLISM | Details ... | 65 | 0.29 | 1.30 | 0.109 | 0.223 | 0.912 | 957 | tags=40%, list=18%, signal=48% |
| 21 | HALLMARK_P53_PATHWAY | Details ... | 91 | 0.27 | 1.29 | 0.114 | 0.221 | 0.919 | 544 | tags=18%, list=10%, signal=19% |
| 22 | HALLMARK_CHOLESTEROL_HOMEOSTASIS | Details ... | 30 | 0.35 | 1.27 | 0.172 | 0.234 | 0.937 | 396 | tags=23%, list=8%, signal=25% |
| 23 | HALLMARK_DNA_REPAIR | Details ... | 68 | 0.27 | 1.27 | 0.139 | 0.227 | 0.943 | 1170 | tags=44%, list=22%, signal=56% |
| 24 | HALLMARK_WNT_BETA_CATENIN_SIGNALING | Details ... | 17 | 0.38 | 1.24 | 0.226 | 0.253 | 0.968 | 307 | tags=18%, list=6%, signal=19% |
| 25 | HALLMARK_UV_RESPONSE_DN | Details ... | 50 | 0.26 | 1.10 | 0.318 | 0.421 | 0.997 | 409 | tags=14%, list=8%, signal=15% |
| 26 | HALLMARK_ANDROGEN_RESPONSE | Details ... | 46 | 0.24 | 1.01 | 0.458 | 0.568 | 1.000 | 838 | tags=28%, list=16%, signal=33% |
| 27 | HALLMARK_XENOBIOTIC_METABOLISM | Details ... | 54 | 0.20 | 0.85 | 0.709 | 0.843 | 1.000 | 787 | tags=22%, list=15%, signal=26% |
| 28 | HALLMARK_PANCREAS_BETA_CELLS | Details ... | 7 | 0.35 | 0.85 | 0.627 | 0.816 | 1.000 | 809 | tags=43%, list=15%, signal=51% |

| | | | | | | | | | | |
|---|---|---|---|---|---|---|---|---|---|---|
| 29 | HALLMARK_NOTCH_SIGNALING | Details ... | 9 | 0.32 | 0.84 | 0.635 | 0.794 | 1.000 | 322 | tags=22%, list=6%, signal=24% |
| 30 | HALLMARK_SPERMATOGENESIS | Details ... | 26 | 0.22 | 0.81 | 0.714 | 0.815 | 1.000 | 127 | tags=8%, list=2%, signal=8% |
| 31 | HALLMARK_PROTEIN_SECRETION | Details ... | 51 | 0.18 | 0.78 | 0.804 | 0.838 | 1.000 | 575 | tags=16%, list=11%, signal=17% |
| 32 | HALLMARK_E2F_TARGETS | Details ... | 83 | 0.16 | 0.77 | 0.873 | 0.826 | 1.000 | 333 | tags=8%, list=6%, signal=9% |
| 33 | HALLMARK_KRAS_SIGNALING_DN | Details ... | 24 | 0.20 | 0.72 | 0.799 | 0.863 | 1.000 | 1256 | tags=33%, list=24%, signal=44% |
